# A structural perspective on the temperature-dependent activity of enzymes

**DOI:** 10.1101/2024.08.23.609221

**Authors:** Matthew J. McLeod, Sarah A. E. Barwell, Todd Holyoak, Robert Edward Thorne

## Abstract

Enzymes are biomolecular catalysts whose activity varies with temperature. Unlike for small-molecule catalysts, the structural ensembles of enzymes can vary substantially with temperature, and it is in general unclear how this modulates the temperature dependence of activity. Here multi-temperature X-ray crystallography was used to record structural changes from −20°C to 40°C for a mesophilic enzyme in complex with inhibitors mimicking substrate-, intermediate-, and product-bound states, representative of major complexes underlying the kinetic constant *k_cat_*. Both inhibitors, substrates and catalytically relevant loop motifs increasingly populate catalytically competent conformations as temperature increases. These changes occur even in temperature ranges where kinetic measurements show roughly linear Arrhenius/Eyring behavior where parameters characterizing the system are assumed to be temperature independent. Simple analysis shows that linear Arrhenius/Eyring behavior can still be observed when the underlying activation energy / enthalpy values vary with temperature, e.g., due to structural changes, and that the underlying thermodynamic parameters can be far from values derived from Arrhenius/Eyring model fits. Our results indicate a critical role for temperature-dependent atomic-resolution structural data in interpreting temperature-dependent kinetic data from enzymatic systems.

**One-Sentence Summary:** Structural data spanning a 60°C temperature range for enzyme complexes mimicking the substrate-, intermediate-, and product-bound states illuminate how small temperature-dependent structural changes may modulate activity and render parameters deduced from Arrhenius/Eyring plots unreliable.

## INTRODUCTION

Chemical reaction rates increase with temperature. For small-molecule catalysts and substrates, increased rates are associated with increased collision velocities and frequencies arising from increased kinetic energy. For enzyme-catalyzed reactions, additional temperature variation in rates may occur due to changes in the conformational ensembles of the enzyme and substrate. Temperature-dependent enzyme reaction rates are typically measured under saturating substrate conditions, where active sites are fully occupied. Turnover ( *k_cat_* ) then reports only on steps after formation of the enzyme-substrate (ES) complex and is independent of collision frequency, and effects of increased kinetic energy will only be observed for processes that occur after formation of ES through to product release and regeneration of the free enzyme (E + P).

Despite these and other significant mechanistic differences, analysis of *k_cat_* as a function of temperature for enzymatic systems is typically based on Arrhenius or Eyring-Polyani (E-P) formalisms, which were derived from small molecule studies and assume the underlying thermodynamic parameters ( Δ*H*, Δ*S* ) are temperature-independent. However, enzymatic Arrhenius/Eyring plots can be more complex than their small molecule counterparts. For example, they may be linear at lower temperatures and then curve downward at elevated temperatures, with rates *k_cat_* rapidly decreasing above a temperature *T_opt_*. This downward curvature is most often attributed to unfolding, and but may be observed even at temperatures well below the unfolding temperature.^(for ex. see 1–6)^

Three models have been proposed to describe this latter behavior: Macromolecular Rate Theory^1^, the Equilibrium Model^7^ and a model described by Roy, Schopf and Warshel^6^. Macromolecular Rate Theory (MMRT) suggests that the temperature dependence of enzymatic rates is controlled by the heat capacity change between two states that determine activity.^1,8–11^ Rigidification of the enzyme ensemble as it traverses the conformationally restricted transition state results in a negative heat capacity difference (Δ*C_p_* ‡) and downward curvature of the Arrhenius plot after *T_opt_*.^1^ The Equilibrium Model suggests that downward curvature is due to a shift of the equilibrium defining an enzyme’s conformational ensemble toward increasing population of inactive states as temperature is increased, prior to denaturation.^7^ The third model suggests that the more polar GS is more conformationally restricted than the less polar TS.^6^ As temperature increases, there is a loosening of the GS that increases its entropy, leading to a decrease in Δ*S*^‡^, an increase in Δ*G*^‡^ and downward curvature. The first two of these models have been the most discussed, with evidence from kinetic measurements, molecular dynamics simulations, theoretical analysis, and model fitting providing support for each. Testing these models requires observation of functionally relevant changes in conformational ensembles along the reaction coordinate as temperature is changed. More generally, such observations are required to understand observed rate-temperature relationships.

A powerful approach to probing temperature-dependent enzyme structure-activity connection is to determine atomic/near atomic resolution X-ray crystallographic structures over a wide temperature range where enzymes exhibit activity. Ideally, the temperature range should span the lower biological limit of roughly −20°C to the denaturation/unfolding temperature,^15^ and the crystal form should allow sufficient conformational flexibility to preserve activity. Robust methods for collecting high-quality structural data from “native”, cryoprotectant-free crystals that maintain liquid internal solvent at temperatures down to ∼200 K (−73 °C) have been established.^16,17^ Methods and hardware for collecting data with minimal crystal dehydration or degradation at up to ∼90°C have also been demonstrated.^18^ This ∼160°C data collection temperature range, feasible at high-resolution with X-ray crystallography, is currently difficult to match using any other structural probe. Despite this, only a handful of multi-temperature crystallography studies spanning a mechanistically useful temperature range have been reported.^19–26^ A recent publication identified only 11 published crystallographic structures at temperatures above 37°C.^27^

Here we performed crystallography at −20, 0, 20, and 40°C to probe a mesophilic, GTP-dependent phosphoenolpyruvate carboxykinase from rat cytosol (rcPEPCK) that retains activity in its crystalline form. To observe changes in structure related to *k_cat_*, we used complexes representing three states on the reaction coordinate that comprise *k_cat_* (**Scheme 1**).^28,29^

**Scheme 1:**
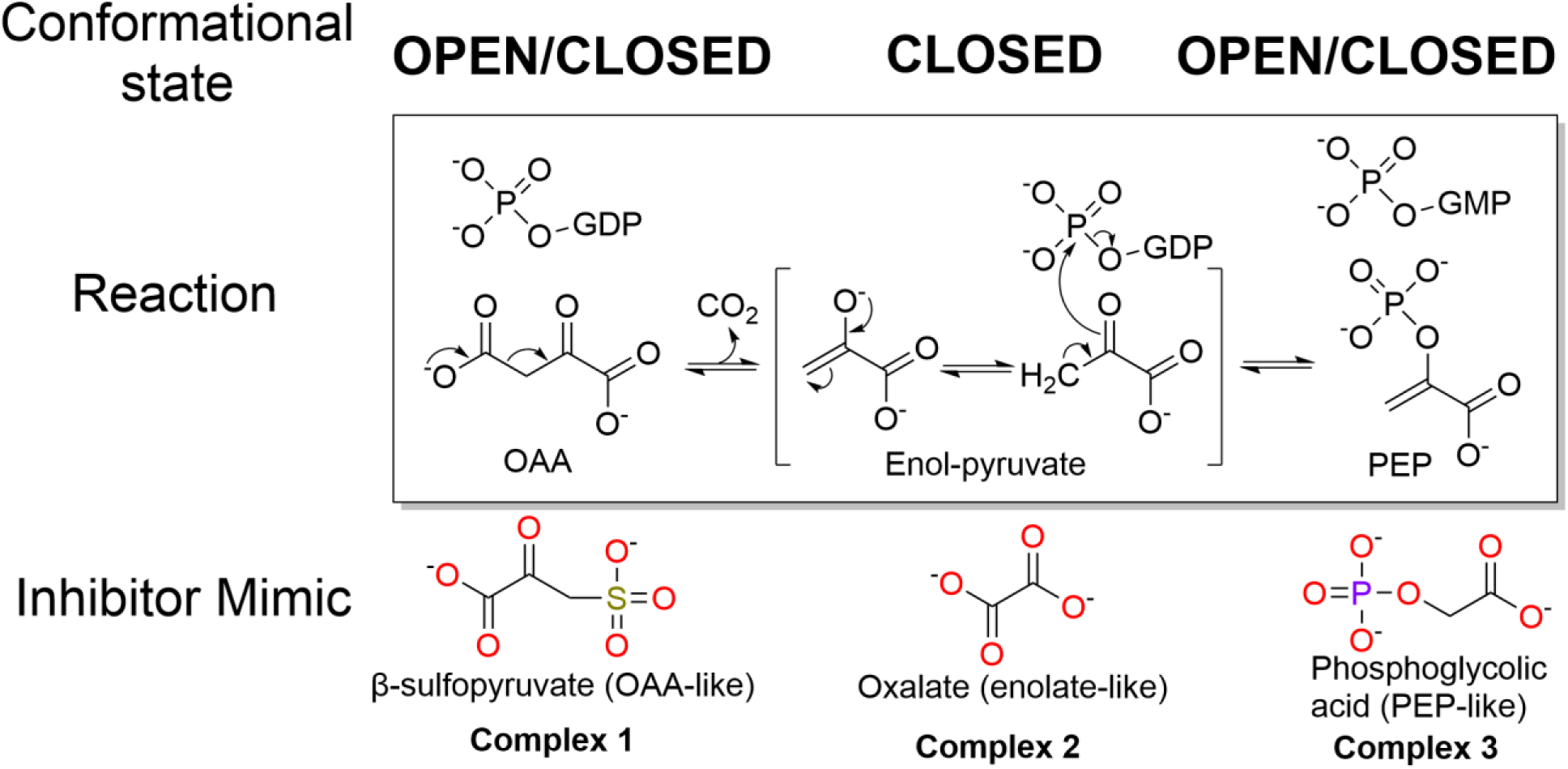
Rat cytosolic PEPCK reaction mechanism and inhibitor complexes.

rcPEPCK has been extensively studied both kinetically and structurally, revealing a clear picture of molecular events leading to activity, allowing temperature-dependent changes to be assessed for their impact on activity.^28–34^ PEPCK is a metabolic enzyme that interconverts oxaloacetic acid (OAA) and phosphoenolpyruvate (PEP) using a metal-cofactor involved in binding and catalysis (M1, typically Mn^2+^ - **Fig. S1**), a second nucleotide-associated metal (M2, typically Mg^2+^), and a phosphoryl donor (either GTP, ATP, or PP_i_ depending on PEPCK class).^35^ With respect to the reversible reaction, the data are consistent with a stepwise mechanism where, in the direction of PEP synthesis (OAA→PEP), the reaction is initiated by the decarboxylation of OAA creating an enol-pyruvate intermediate. This intermediate is subsequently phosphorylated by the phosphoryl donor producing PEP (**Scheme 1**).^(reviewed in 35)^ Substrate binding and activity have been shown to be coupled to several dynamic transitions in the enzyme, including a global closure via rotation of the N- and C-terminal domains that reduces the active site cavity’s total volume, as well as loop and residue rearrangements.^32^ More specifically it has been shown that the R-(substrate-binding) and P-(nucleotide-binding) loops undergo disorder-to-order transitions upon binding, aiding in orienting the substrate and nucleotide appropriately.^32^ The Ω-loop, an active site lid, undergoes a similar essential disorder-order transition as it folds and closes over the active site and is held in place by the adjacent R-loop after the enzyme has undergone substrate-induced global closure and active site remodeling (**Fig. 1**).^31,33,36^ Lid closure has been shown to be essential to PEPCK function as the lid holds the active site and substrates in a catalytically competent state and protects the enol-pyruvate intermediate from solvent-mediated protonation and the non-productive formation of pyruvate.^36^

**Fig. 1.**
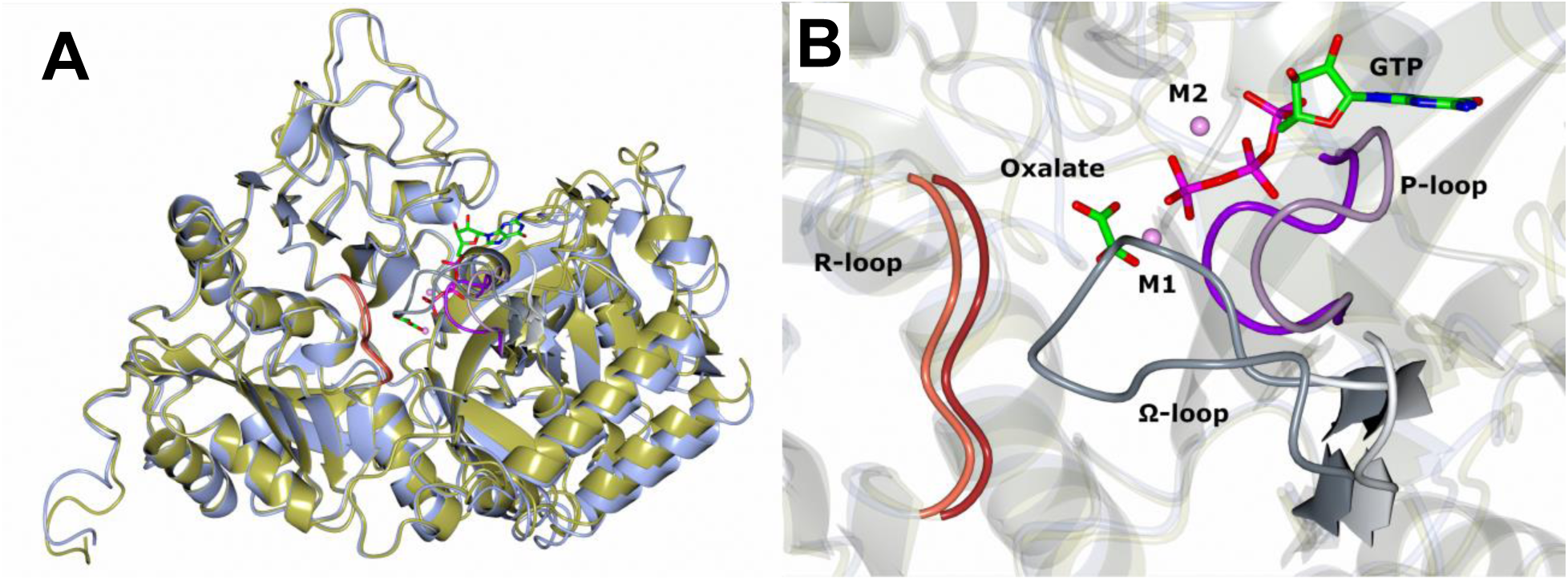
Open and closed conformations of rcPEPCK along the reaction coordinate. Superposition of the open and closed structures (N-terminal domain residues 3-250) of rcPEPCK, indicating conformational changes of A) the global structure and B) the active site. Holo rcPEPCK (open) is shown in steel blue (PDB 2QEW), and its Ω-loop in light grey, R-loop in coral, and P-loop in lilac. Oxalate- (enol-pyruvate intermediate mimic) GTP bound rcPEPCK (closed) is shown in gold (PDB 3DT2) and its Ω-loop in grey, R-loop in firebrick red, and P-loop in purple. Oxalate and GTP are shown bound to the active site and atoms are colored by type (red – oxygen, blue – nitrogen, green – carbon, purple – phosphorous, pink – manganese).

Our temperature-dependent structural characterization of rcPEPCK, as well as other previous works indicating structural changes with varying temperature^19,20,22,37,38^, prompted us to re-evaluate assumptions made in applying Arrhenius / Eyring-Polanyi (E-P) models to temperature-dependent enzyme kinetic data. Although Arrhenius/E-P plots provide a qualitative descriptor of an enzyme’s free-energy landscape, combined multi-temperature structural and kinetic/biochemical measurements are required for quantitative and mechanistic insight.

## RESULTS

### Multi-temperature kinetics

An Eyring-Polyani (E-P) plot of *k_cat_* for rcPEPCK in the reverse, PEP→OAA, direction (**Fig. 2**, raw data in **Data S1**) showed a decreasing slope with increasing temperatures above ∼25°C (**SI Data S2**), with a maximum rate at ∼50°C ( *T_opt_* ) and a loss of activity at higher temperatures. In the presence of viscogen (glycerol) at constant viscosity (2.4 cp/mPa), *k_cat_* decreased at temperatures above ∼50°C but was unaffected at lower temperatures. *k_cat_* / *K_M_* for the PEP→OAA reaction was determined at four temperatures between 15°C and 55°C (**SI Table S1)** but the data are not sufficient to assess the functional variation with temperature. *k_cat_* for the OAA→PEP direction could not be reliably determined above 35°C as the non-enzymatic rate of metal-catalyzed decarboxylation of OAA became significant. An Arrhenius plot of the available data (7-37°C) appeared roughly linear (data not presented).

**Fig. 2.**
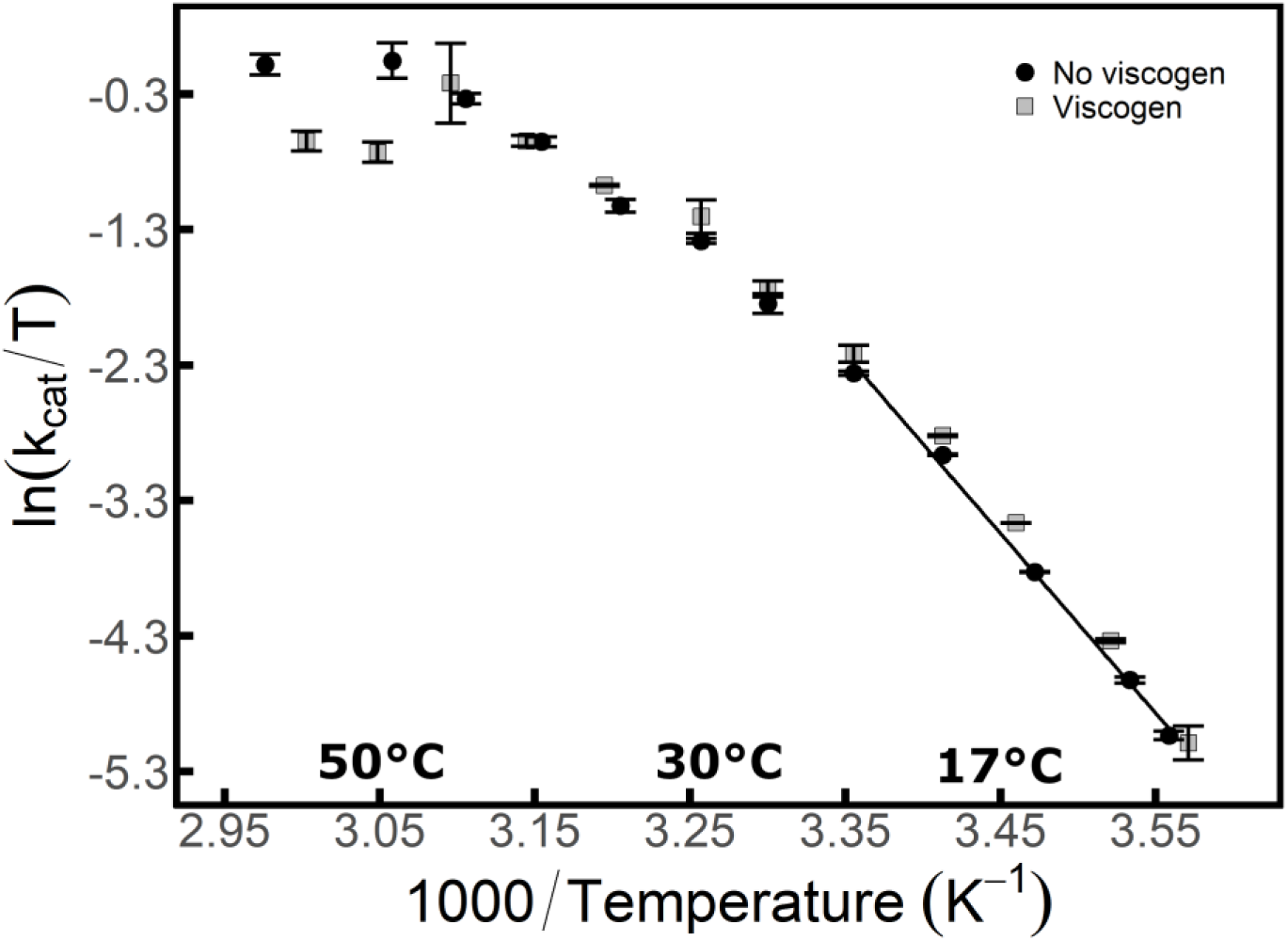
Eyring-Polanyi plot for rcPEPCK mediated carboxylation of PEP. Reaction rate vs. temperature at saturating concentrations of substrate without (black circle) and with viscogen (grey squares). Linear fit corresponds to 8-25°C temperature range. Error bars (standard error) may be hidden within the data point.

### Multi-temperature crystallography

While all three complexes examined here were inhibited complexes using mimics for the OAA/PEP substrates, structural evidence using authentic substrates with WT and mutant forms of the enzyme suggest that the conformational states sampled in these complexes faithfully reproduce those of the catalytically competent enzyme-substrate complexes (**Fig. 1 and SI Fig. S2**).^28,29,34,39^ In the oxalate(OX)-GTP and phosphoglycolic acid (PGA)-GDP complexes CO_2_ is not present. Therefore, the PGA-GDP complex represents a partially occupied encounter/product complex (prior to CO_2_ binding / after CO_2_ is released) and the OX-GTP complex represents the intermediate state which may or may not have CO_2_ present in the authentic enolate-GTP complex but is still conformationally closed. Therefore, both these complexes represent meaningful states along the catalytic trajectory.^28,39^

#### rcPEPCK β-sulfopyruvate (βSP)-GTP complex (**Scheme 1**, Complex 1)

Both βSP and GTP were fully occupied in the active site at all temperatures. For GTP, the electron density indicated two resolvable conformers between −20°C and 20°C (**Fig. 3**). One conformer was modeled in a fixed position that has been clearly resolved in the open PEPCK-GTP complex at cryogenic temperatures, and is deemed incompetent for phosphoryl transfer (**IC** in **Fig. 3**).^29^ The electron density suggests the other conformer changes pose with temperature, and at 40 °C adopts a conformation that has been observed in cryogenic structures of the closed, oxalate-GTP complex, deemed competent for phosphoryl transfer (**C** in **Fig. 3**).^28,30^ At 40°C, there is only evidence for the closed oxalate-GTP complex conformer. The conformation change of GTP occurs by a rotation of both the α-phosphate and ribose out of the plane of the triphosphate moiety. Occupancy refinements, when the B-factors are fixed to the proteins average B-factor, shows the fixed PEPCK-GTP complex conformer is depopulated while the rotating oxalate-GTP like conformer is populated with temperature (**Fig. 3**). The occupancies for IC:C conformers are 72%:28% at −20°C, 52%:48% at 0°C, 39%:61% at 20°C, and 0%:100% at 40°C. The change in nucleotide conformation was coupled to movement of the P-loop as it shifted towards the M1 metal with increasing temperature - a process previously associated with enzyme/lid closure in cryogenic structures (**SI Fig. S3**).^28,29,40^ These adjustments lead to various increases and decreases in interatomic distances between GTP and contacting residues (**SI Fig. S4**). Electron density in the ordered Ω-loop conformation was observed to increase between −20°C and 0°C, remained nearly constant between 0°C and 20°C, and diminished at 40°C (where overall resolution was worse), hinting at possible disordering of the lid at 40°C and above (**Fig 4, Table S2,** and **Fig. S5** for βSP-GTP Ω-loop electron density at lower contour). Over the −20°C to 20°C temperature range where data set resolution was nearly constant, the normalized B-factors (B-factor_res_-B-factor_protein_) of residues 94-134 and 228-251, increased with increasing temperature while those of residues 59-93, 255-303, 413-420, 535-557, and 579-590 decreased (**SI Data S3**). The normalized B-factors for the Ω-loop are consistent with increased order with increasing temperature between −20°C to 20°C evident in the electron density maps.

**Fig. 3.**
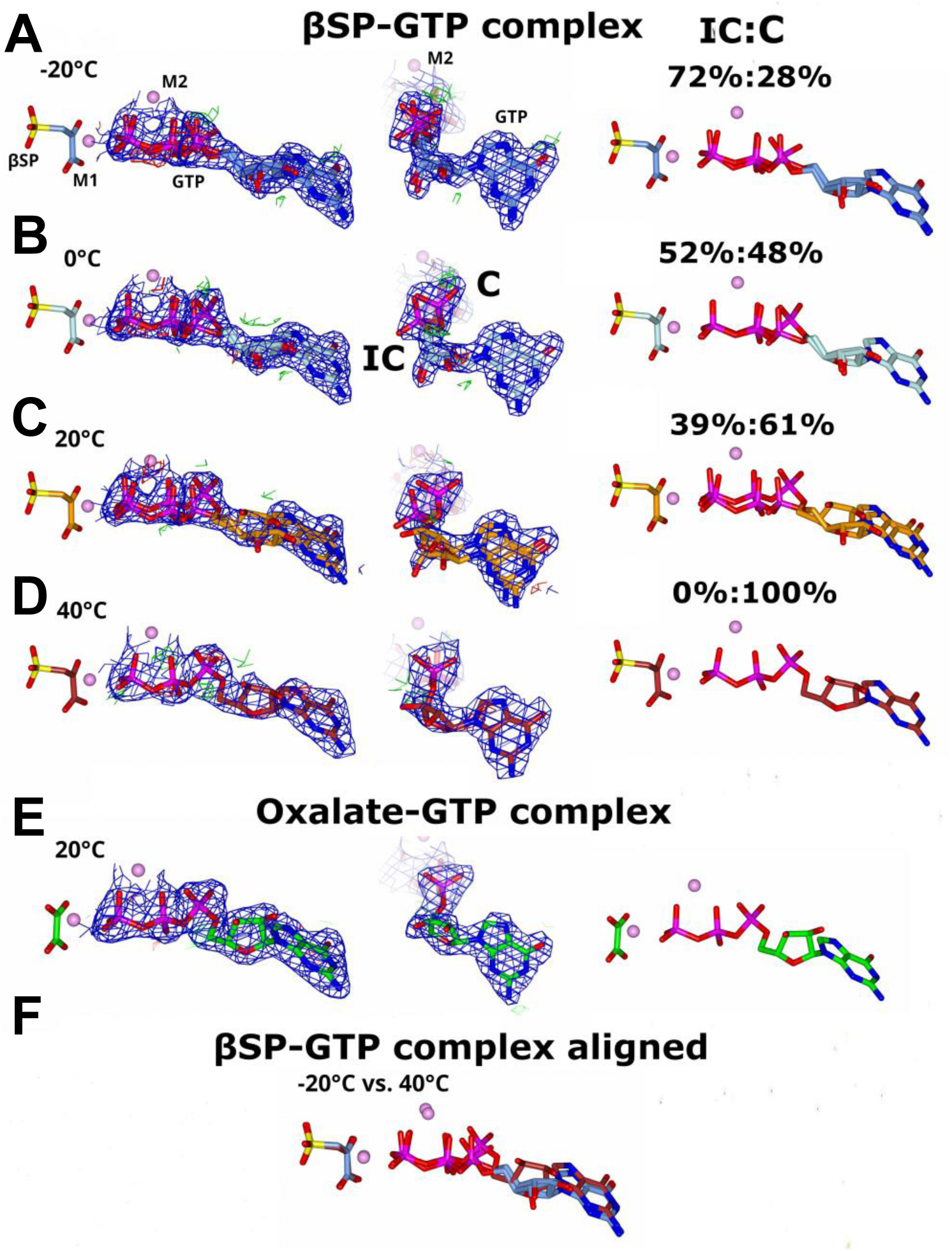
Conformational change toward intermediate state with increasing temperature in the βSP-GTP (forward reaction GS) complex. Structures determined at A) −20°C (carbon – ice blue), B) 0°C (light blue), C) 20°C (orange), and D) 40°C (firebrick red) show a shift in GTP conformation as temperature is increased from the fixed, incompetent position to the rotated, competent one (IC and C respectively). The middle panel shows the same structure rotated 90° (looking in-line with nucleotide) to show conformational changes of the α-phosphate. The right panel shows the modeling without electron density with the refined occupancy values for each conformer. Specifically, the α-phosphate and ribose rotate away from the in-line plane of the triphosphate tail. At 40°C, the conformation of the nucleotide is almost identical to E) the 20°C intermediate complex (green – oxalate-GTP). F) Alignment of −20°C and 40°C structures showing the extent of the conformational change. All other atoms are colored by atom-type. Both 2F_o_−F_c_ (blue – 1σ) and F_o_−F_c_ (green and red – 3σ) electron density maps are shown.

**Fig. 4.**
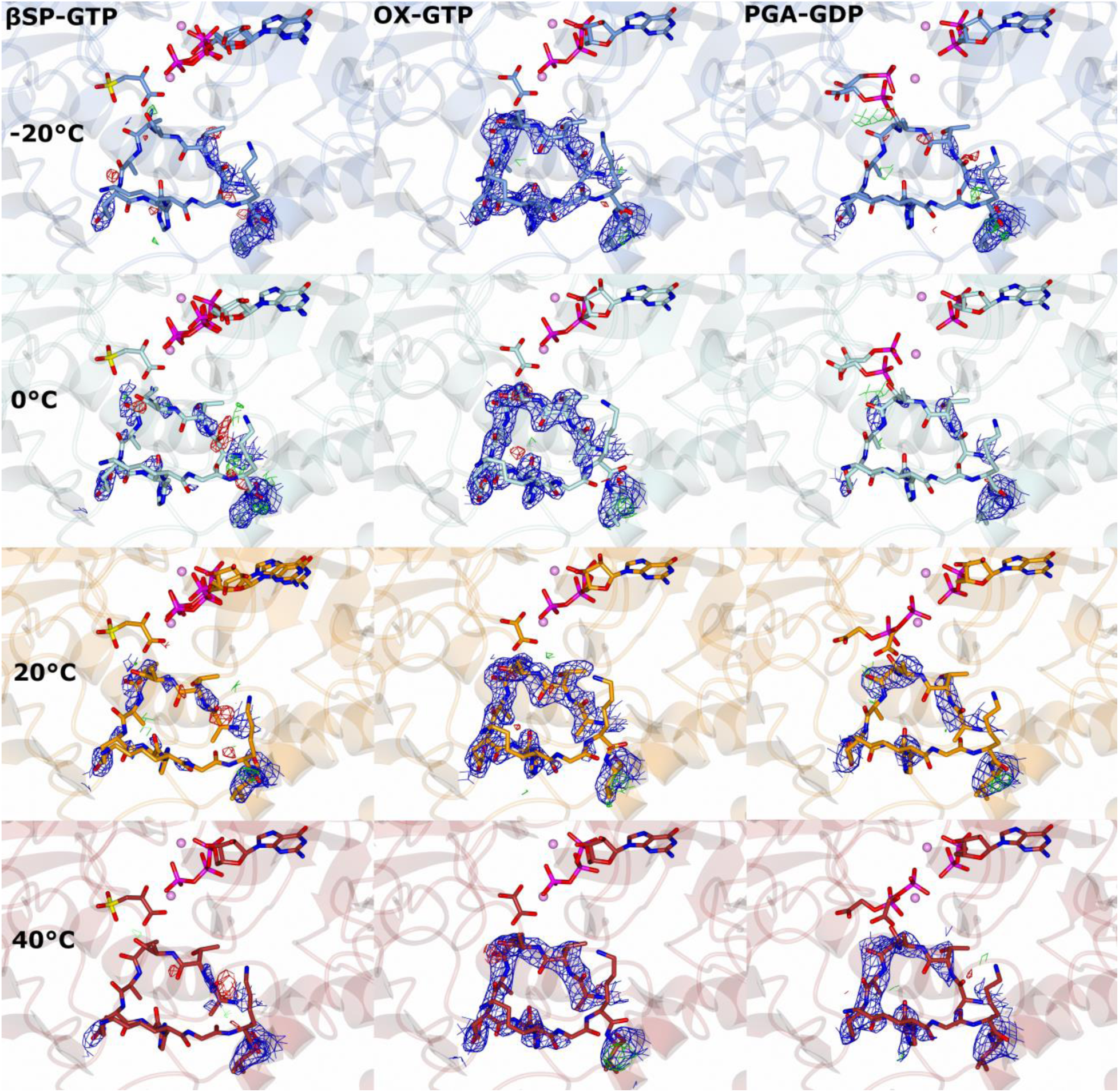
Changes in Ω-loop closed conformation occupancy. In each complex, the Ω-loop is modelled in the closed conformation with an occupancy of 1.0. The 2F_o_−F_c_ (blue) and F_o_−F_c_ (red/green) maps resulting from refinement are contoured to 1σ and 3σ, respectively. In the βSP-GTP complex, the closed-conformation occupancy of the Ω-loop increases from −20°C to 20°C, and then shows weak evidence of opening/disordering by 40°C (**Fig. S4**). In the oxalate-GTP complex, the Ω-loop is mostly closed at all temperatures. In the PGA-GDP complex, the closed conformation occupancy of the Ω-loop increases with increasing temperature.

#### rcPEPCK oxalate (OX)-GTP complex (enol-pyruvate-GTP) (**Scheme 1**, Complex 2)

In contrast to the temperature-dependent changes noted for the βSP-GTP complex **(Complex 1)**, a single dominant/resolvable conformer of oxalate and GTP was modeled at all temperatures. The solvent-exposed O2’ of the ribose sugar (**SI Fig. S6**) and adjacent atoms showed a significant increase in B-factor with increasing temperature, consistent with this region of the nucleotide becoming disordered rather than populating a new conformation. The lack of movement in the triphosphate portion is correlated with a single dominant/resolvable conformation of the P-loop at all temperatures (**SI Fig. S3**). Strong electron density for the Ω-loop was present at all temperatures (**Fig. 4**) although a slightly larger B-factor may suggest increased disorder at 40°C (**SI Table S2**). Consistent with prior cryogenic structures, at all temperatures the OX-GTP complexes had a PEG molecule bound in a cavity between the N- and C-terminal domains of PEPCK; a loop formed by residues 149-154 changed position to accommodate the PEG. In both the βSP-GTP and PGA-GDP complexes, the cavity region was partially disordered but in the dominant/resolved conformation the loop closed off the cavity to prevent PEG binding (**Fig. S7**). B-factor analysis indicates some minor temperature dependencies in peripheral regions of the enzyme in this complex. (**SI Data S3**). Residues 95-125 show increased disorder while residues 265-300 show increased order with increased temperature.

#### rcPEPCK phosphoglycolic acid (PGA)-GDP complex (PEP-GDP) (**Scheme 1**, Complex 3)

Like the OX-GTP complex, in the PGA-GDP complex at all temperatures, GDP was fully occupied in the active site and single dominant/resolved conformations of the P-loop and nucleotide were observed (**SI Fig. S3**). The O3’ and other ribose atoms exhibited local temperature-dependent disorder (**SI Fig. S6**), as in the OX-GTP complex. PGA, on the other hand, exhibited significant conformational change with temperature (**Fig. 5** and **Fig. S2 & 8**). PGA (and PEP) are known to adopt three discrete conformations.^28,34,39^ Two of these exhibit second sphere coordination to the M1 cation and positions the phosphate at too great a distance from the β-phosphate of GDP for phosphoryl transfer to occur, and are thus deemed catalytically incompetent (IC1, IC2) (**Fig. 5** and **Fig. S2 & 8**).^39^ In the third conformation, PGA (PEP) directly coordinates to the M1 metal, and the phosphate is positioned in the same location as the γ-phosphate of GTP that is unoccupied when GDP is bound, and is therefore the catalytically competent conformation (C1, **Fig. 5** and **Fig. S2 & 8**).^28,34^ At −20°C, IC1 and IC2 were fully occupied with a distribution of 40%:60%:0% (IC1:IC2:C1), and shifted to a 50%:50%:0% distribution at 0°C. At 20°C, C1 became populated with a 0%:55%:45% distribution, and at 40°C the shift toward the competent state was nearly complete with a 0%:25%:75% distribution. Concomitant with this binding mode change, Y235 shifted from a buried conformation when the incompetent conformations were dominant and rotated toward the active site as the competent conformation was populated (**Fig. 5D)**. The Ω-loop made a similar shift: it largely occupied the open, disordered state at −20°C, and became increasingly ordered and closed as temperature increased (**Fig. 4**). The only indication of temperature dependency from normalized B-factors was the increase in order of the Ω-loop as temperature increased (**SI Data S3**).

**Fig. 5.**
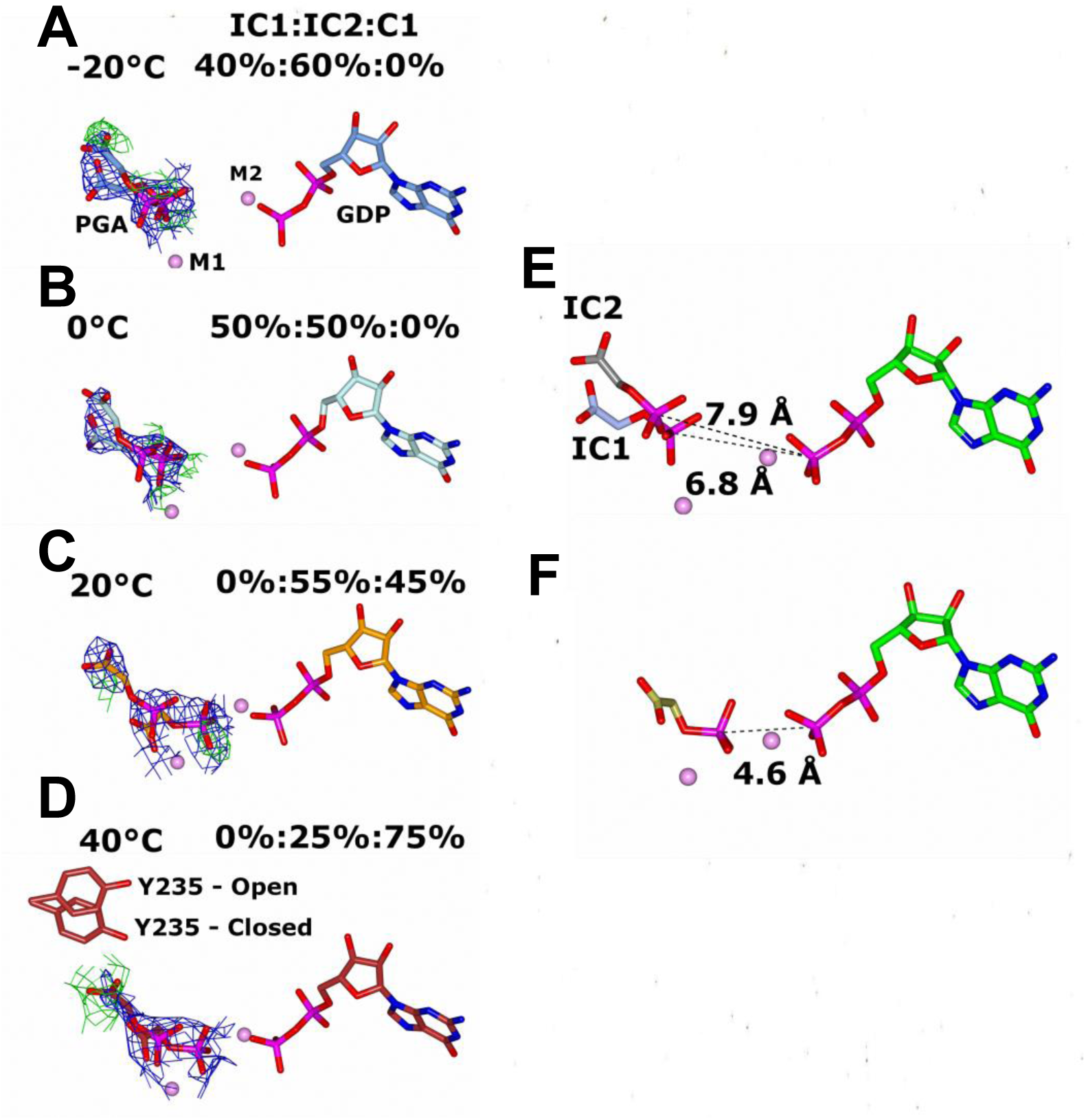
Electron density maps for PGA indicate substrate conformational change with temperature. Model and electron density difference maps at A) −20°C, B) 0°C, C) 20°C, D) 40°C. PGA was observed in three distinct conformations: E) two incompetent outer shell states (IC1 and IC2; PGA-GDP P-P interatomic distances of 7.94 and 6.84 Å) and F) one competent state coordinating directly to the M1 metal (C1; P-P interatomic distance 4.84 Å) (**Fig. S1)**. As temperature increased, the relative population of the competent conformation increased. Each occupancy was manually refined to minimize the difference density present and the resultant occupancies are labeled (IC1:IC2:C1) for each panel. Y235 is shown in D) indicating the coupled conformational change of the active site with PGA/PEP location. The final 2F_o_-F_c_ maps (blue) are contoured to 1σ, and F_o_-F_c_ (red/green) maps are contoured to 3σ. Positive difference density in D) surrounding IC2 was unchanging as its occupancy is increased.

## DISCUSSION

The significance of crystallographic findings depends on constraints imposed by the crystal lattice and how these modify the ensemble of accessible conformational states relative to those of the enzyme in solution. Prior characterization of rcPEPCK crystals has found that the lattice allows multiple conformational states to be observed. For example, all catalytically relevant domain and loop motions including the transition between open and closed states upon soaking with substrates or inhibitor mimics are allowed. The rich history of structure-function investigations of rcPEPCK provides a basis for interpreting temperature-induced changes.^28–35,39,41–44^ Furthermore, a simplified approach to time-resolved crystallography has recently revealed the hypothesized active conformation of PEP (*vida infra,* competent 1 (C1) conformation) generated from OAA and GTP, confirming that rcPEPCK is active *in crystallo* and that all dynamic modes necessary for activity are accessible (**SI Fig. S2**).^34^

### Temperature increases global B-factors but has variable and complex-dependent local effects

Nearly all previous “more than one” multi-temperature crystallographic studies have focused on comparison of room temperature ∼298 K (25°C) and cryogenic temperature ∼100 K (−173 °C) structures. These studies have revealed significant changes in the structural ensemble associated with cryocooling and indicate the importance of measuring structures at physiological temperatures in assessing structure-function relationships.^21,23–25,45–48^ Data collected at temperatures above 100 K and below ∼220 K (−53°C) (near and below the protein-solvent dynamical (or glass) transition, where most functionally salient protein and solvent dynamics are absent) usually add limited information of relevance to understanding mechanism beyond that available at 100 K.^20^ Where crystallographic data at several temperatures above the dynamic transition has been collected, only the apo / holo form and one additional complex have been examined, rather than a larger set of complexes representing states along the reaction coordinate.^19,20,22,26,49^ Multi-temperature cryoEM studies, where grid-containing samples are prepared at different temperatures immediately prior to cryocooling, have examined two systems and revealed domain-scale changes and changes in the ligand pose.^37,38^ For one system only the apo complex and the ternary “transition state” like complex^37^ were examined, at 6 temperatures between 4°C and 70°C and resolutions between 2.1 and 3.0 Å.^37^ For the other system, wild-type and mutant versions of a single complex were examined at temperatures of 4, 37 and 42 °C and resolutions of 4.1 - 5.2 Å. Unlike in multi-temperature crystallography where the target temperature is maintained during data collection, multi-temperature cryoEM requires freeze-trapping which perturbs the conformational ensemble. Even though cooling rates of thin-film cryoEM samples are one to two orders of magnitude larger than can be achieved with microcrystals, substantial relaxation of side chains and smaller moieties is still expected, and sample precooling in cold gas present immediately above the liquid ethane introduces uncertainty in the trapped temperature.

Previous multi-temperature structural studies revealed two general phenomena which the present data support. First, global average B-factors increase as temperature increases, primarily reflecting increased thermal fluctuations rather than increased static disorder.^20–22,50^ Second, the effects of temperature on the dynamic behavior of local structural elements such as residues and loops depend on the local context of neighboring residues, charge distributions/pKas, solvation, steric interactions, etc. Both increases and decreases in order with increasing temperature are observed.^19–21,23^

Here, the global B-factor for each of three sampled rcPEPCK complexes comprising *k_cat_* increased with increasing temperature (**SI Data S4**). Each complex has a unique temperature dependence, with local disorder either increased or decreased depending on the region examined (**SI Data S3**). The intermediate OX-GTP complex was the most temperature agnostic, as the global B-factor showed only a small change and the Ω-loop and substrates did not show clear changes in position or occupancy with temperature. In contrast, both the forward (βSP-GTP) and reverse (PGA-GDP) GS complexes showed an increase in Ω-loop (active-site lid) order with increasing temperature, with additional ordering in the PGA-GDP complex but increased disorder in the βSP-GTP complex observed at 40°C (**Fig. 4, Table S2** and **Fig. S3**). There was also an increase in the normalized B-factor (**Table S2**) of the Ω-loop of the OX-GTP complex at 40°C, but this does not translate to significantly more disorder (**Fig. 4**). Coincident with increasing lid order, the P-loop and nucleotide conformations in the βSP-GTP complex change (**Fig. 3 and Fig. S3)**. At 40°C, GTP adopts an eclipsed, competent conformation; the γ-phosphate becomes a better leaving group, thus promoting phosphoryl transfer and turnover.^28,40^ This eclipsed geometry is similar to that observed in the intermediate state (OX-GTP) (**Fig. 3).** Although this conformational change which aids phosphoryl transfer (second chemical step) occurs in the βSP-GTP complex, before decarboxylation (the first), it will still result in an increase *k_cat_*. At high temperatures, phosphoryl transfer can immediately follow decarboxylation because GTP is already in the eclipsed geometry. In contrast, at low temperatures, after decarboxylation GTP must undergo an energetically costly conformational change to the eclipsed state before phosphoryl transfer can occur. This additional conformational change results in an increased time for turnover. In the PGA-GDP complex, PGA shifted from predominantly populating the two incompetent conformations (IC1 and IC2) below 20°C toward increasing occupancy of the competent state at 20 and 40°C — a conformational shift that is required for phosphoryl transfer (C1) (**Fig. 5** and **SI Fig. S8**).

### Multi-temperature crystallography reveals structural changes influencing kinetic parameters

The observed structural changes are directly supported by previous structure-function studies indicating similar changes when rcPEPCK traverses its reaction coordinate.^28–31^ Together with results from previous cryoEM studies^37,38^, the present results sampling enzyme-ligand complexes corresponding to known states comprising *k_cat_* suggest that the equilibrium between conformational states shifts so that the population of active states increases with increasing temperature up until some maximum, *T_opt_*.

This increase in active state population likely contributes to the observed increase in *k_cat_* with temperature, as this parameter reflects the energetic barriers of all events on the reaction coordinate following the formation of the enzyme-substrate complex.^51^ If one of these events has a *significantly* slower rate than the others, *k_cat_* will be *mostly* controlled by that event and following events; this is often simplified by assuming that *k_cat_* represents this one event. However, in measurements over a large temperature range (e.g., from ∼280 K to 340 K), temperature-dependent structural changes may differentially affect the individual microscopic steps and their relative contributions to *k_cat_*, as seen in the different temperature variations of the Ω-loop in each sampled complex. Here, significant differences in the structural data in response to changing temperature of inhibitor complexes representing three meaningful states on the reaction coordinate are observed. These data indicate changes that may plausibly impact rate-determining step(s) at low and high temperatures.Alternative possibilities that could impact rate-determining steps include changes to the dominant reaction mechanism itself, perhaps involving an increase in occupancy of competent conformers with occupancies too low to be discerned in our maps; and changes in the TS structure with temperature.

A previous mutational and kinetic study found that rcPEPCK is at least partially rate-limited by phosphoryl transfer (at ambient temperature).^33^ The present structural data support this conclusion, as the structural changes observed (eclipsed phosphate geometry of GTP, PGA conformation changing to coordinate with M1, lid closure allowing for chemistry) with increasing temperature all lead to a shift in conformational state that is more favorable for phosphoryl transfer.

The intermediate state, represented here by the OX-GTP complex, is resistant to temperature-induced changes. This is not surprising, as the conformational ensemble of this complex must be tightly constrained for the reaction to proceed. Prior work demonstrated that under conditions where the Ω-loop opens prior to phosphorylation of the enolate intermediate, solvent can protonate the intermediate to form a side product, pyruvate.^31^ PEPCK likely evolved to ensure robust loop closure throughout the chemical steps essential to maximize the enzyme’s efficacy, consistent with the conformational stability observed for the OX-GTP complex. The induction of the closed conformation in the OX-GTP complex is likely aided by rotation of Y235, as all other active site residues have largely the same positions in the βSP-GTP and OX-GTP complexes (**Fig. S8**)^28^, by a redistribution of interaction distances at the active site (and beyond), and by the physicochemical properties of the ligand itself.

### Increased local disorder in the product-complex suggests a new rate-limiting step at high temperatures

Arrhenius / E-P plots of *k_cat_* (*T*) for rcPEPCK exhibit non-linear behavior at high temperature, with negative curvature leading to a maximum value for *k_cat_* (*T*) at a *T_opt_* of ∼50°C and slope inversion at temperatures above *T_opt_.* As mentioned previously, this non-Arrhenius behavior near *T_opt_* is common in enzyme systems, and recent efforts have attempted to create a generalizable model describing it.^1–3,5,6,13,14,52,53^ Based upon these efforts, negative curvature at temperatures near *T_opt_* may be attributed to 1) a change in mechanism, 2) an increase in the barrier that limits rates at high temperatures, or 3) a new barrier along the reaction coordinate becoming dominant. For rcPEPCK, significant downward curvature, starting near ∼40°C, coincides with *k_cat_* exhibiting sensitivity to solvent viscosity (∼50°C). A possible interpretation is that a new, non-chemical rate-limiting step has emerged (**Fig. 2**). Structures at 40 °C provide weak evidence of a shift to product release as the rate limiting step, which would be consistent with a high-temperature viscosity dependence of *k_cat_*. Under these assumptions, we speculate that lid ordering prior to chemistry may then become increasingly unfavorable, and this dynamic (and presumably solvent friction-sensitive) step could become rate limiting.

Attempts to obtain high-resolution crystallographic data at temperatures above 40°C were not successful due to crystal instability. However, structures at 40°C provide weak evidence that a shift to product release as the rate limiting step may play a role in the high-temperature viscosity dependence of *k_cat_*. For the PEP→OAA reaction direction, at 40°C the Ω-loop in the βSP-GTP (product) complex shows hints of increased disorder: a loss of ordered density (**Fig. 4 and Fig. S5**) in the loop and an increase in B-factor that is larger than expected based solely on the lower resolution of the 40°C data set (**Table S2**). Disordering of the Ω-loop should aid in product release thus increasing *k_cat_*. In the PGA-GDP (substrate) complex, the lid remains closed at 40°C. Given that the Ω-loop has no direct interaction with either the product or substrate complex, it seems plausible that, like the βSP-GTP complex, disorder in the lid may develop above 40°C. Under these assumptions, we speculate that lid ordering prior to chemistry may then become increasingly unfavorable, and this dynamic (and presumably solvent friction-sensitive) step could become rate limiting. Alternatively, new high-temperature conformers may arise which can specifically bind glycerol. This binding may then increase the fraction of inactive conformers. These hypotheses can be tested with molecular dynamics simulations.

### Linear Arrhenius plots hide a changing free-energy landscape that is revealed through multi-temperature crystallography

The temperature dependent activity of related enzymes (e.g., two mutants) is often compared using Arrhenius or Eyring plots. Activation / thermodynamic parameters are extracted from the slope and *y*-intercept of linear fits. These plots typically report a modest number of data points, each with significant experimental uncertainty, acquired over a limited temperature range. Uncertainty in the slope of such fits arises both from measurement uncertainties and from using a linear fit when the underlying variation may be nonlinear. Uncertainties in the *y*-intercept are particularly large, as a fit to data spanning, e.g., 0.003 K^-1^ <1/T<0.0037 K^-1^ must be extrapolated over more than four times that interval to 1/T=0.^54^

An implicit assumption in applying the Arrhenius and Eyring models is that, when *k_cat_* (or *k_cat_* / *T* varies linearly with temperature over some range, as is the case at low temperature for rcPEPCK, the thermodynamic parameters describing the system are temperature-independent. Thus, a single “apparent” activation entropy Δ*H^rxn^*^,*app*^ and enthalpy Δ*H^rxn^*^,*app*^ (delineated from the true values Δ*S^rxn^* and Δ*H^rxn^*)) are assumed. This may be a reasonable assumption in small-molecule reaction kinetics. However, hydrophobic interactions, pK_a_ values, and other properties/interactions of relevance to protein structure and dynamics are temperature dependent.

As is clear from this and previous multi-temperature structural studies, enzyme structure is temperature dependent. Here we have observed functionally relevant changes in inhibitor-bound active site structures mimicking the complexes sampled by with *k_cat_* even in temperature ranges where the corresponding kinetics plots appear to be strictly linear (**SI Data S2**). Based on these observations for enzyme systems, there is no reason to expect the underlying thermodynamic parameters to remain constant over the temperature ranges typically probed in kinetics experiments.

### Interpretation of kinetics measurements

As will be discussed in more detail elsewhere, a linear E-P fit to the rcPEPCK data (implicitly assuming a temperature-independent free-energy landscape) (**Fig. 2**) between 8°C (281 K) and 25°C (298 K) gives Δ*H ^rxn^*^,*app*^ = 26.3 ± 0.85 kcal/mol and Δ*S^rxn^*^,*app*^ = 0.037 ± 0.003 kcal/mol•°C (*R*^2^=0.997). Identical fits to the rcPEPCK data can be obtained by assuming that Δ*H^rxn^* at 8°C is 21.30 kcal/mol (19% smaller) and decreases with increasing temperature at a rate of 0.0178 kcal/mol•°C (i.e., by ∼0.25 kcal or ∼1/4 of a hydrogen bond over 14 °C) and a constant Δ*S^rxn^* = 0.019 kcal/mol•°C (48% smaller); or by assuming that Δ*H^rxn^* at 8°C is 16.3 kcal/mol (39% smaller) and decreases with increasing temperature at a rate of 0.035 kcal/mol•°C (i.e., by ∼0.5 kcal or ∼1/2 of a hydrogen bond over 14 °C) and a constant Δ*S^rxn^* = 0.016 kcal/mol•°C (96% smaller). In fact, there is a wide range of physically plausible combinations of linearly varying enthalpies Δ*H^rxn^* (*T*) and fixed entropies Δ*S^rxn^* (or of simultaneously varying enthalpies and entropies) that can recapitulate the E-P fit to the activity data. In other words, the observed slope and intercept of Arrhenius or E-P plots of *k_cat_* (*T*) constrain but do not determine the underlying enthalpies Δ*H^rxn^* (*T*) and entropies Δ*S^rxn^* (*T*) and their temperature dependencies that determine the observed *k_cat_*.

This analysis indicates that small temperature variations of underlying thermodynamic / reaction parameters may, in the absence of near-atomic resolution multi-temperature structural information and/or support from other measurements, make quantitative and even qualitative interpretation of parameters derived from Arrhenius / E-P fits to enzyme kinetic data unreliable, with significant consequences for mechanistic interpretations. Qualitative ordering of, e.g., mutant enzymes based on their Arrhenius / Eyring slopes and intercepts may be unreliable.^48,55–59^ Kinetics data only yields Δ*G^rxn^* (*T*), a composite of temperature-dependence, and unknown enthalpy and entropy.

## CONCLUSION

Stochastic thermal fluctuations of an enzyme-substrate complex drive the crossing of energetic barriers. Unlike in small molecule catalysis, in enzymes the relevant enthalpy and entropy changes must vary with temperature due to an enzyme’s many degrees of freedom and lability, the many steps along the reaction coordinate^60^ contributing to the overall Δ*G^rxn^* (*T*), and the temperature variations of physicochemical interactions that govern enzyme structure and activity. Together, these modulate the temperature-dependent activity that would be otherwise be observed if the enzyme were fully “locked down”, with the loss of activity accompanying high temperature unfolding being the most obvious example. Consequently, analysis of kinetic data based on specific mechanistic assumptions (e.g., a temperature-independent free-energy landscape) may be confounded by gradual variations of underlying parameters ( Δ*H^rxn^* and Δ*S^rxn^*) with temperature.

To better understand the temperature-dependent activity of enzymes, we used multi-temperature crystallography to reveal the structural changes of rat cytosolic PEPCK-inhibitor complexes, an extensively characterized model system. We have identified temperature-dependent changes in the active site and ligand conformations that, based on extensive prior studies, are expected to contribute to the observed increase in *k_cat_* with increasing temperature at temperatures Below *T_opt_*. Generally, as temperature increases there is a shift from less competent to more competent active site / substrate configurations, contributing to the increase in *k_cat_* (and decrease in Δ*G^rxn^* (*T*)) observed over the same temperature range. These structural changes are observed even within temperature intervals where the corresponding E-P plots appear linear. At the highest temperature probed, weak evidence for increased disorder of the active site lid suggests a shift in this equilibrium back toward chemically incompetent enzyme states, that in turn may be the origin of the negative Arrhenius curvature and loss of activity at higher temperatures.

Motivated by these and previous structural observations, it is clear than the assumption of temperature-independent parameters ( *E_a_*, *A* or Δ*H^rxn^*, Δ*S^rxn^* ) when interpreting Arrhenius or Eyring-Polanyi fits will in general be invalid, even when fitting temperature intervals over which plotted data appears linear. As will be shown elsewhere, small temperature variations of *E_a_* and Δ*G^rxn^* will have large and correlated effects on the slope and intercept – on 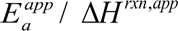 and *A^app^* / Δ*S^rxn,app^*, muddling their mechanistic significance. Effects of these temperature variations may be so large as to render comparisons of fit parameters Δ*H^rxn^*^,*app*^ and Δ*S^rxn,app^* obtained from related enzymes, including engineered enzymes and enzymes adapted to different thermal environments, largely meaningless unless observed differences are supported by complementary measurements. Our results point to a key role for atomic resolution multi-temperature structural studies in illuminating fundamental aspects of structure-function relationships, protein design, and thermal adaptation.

## Acknowledgments

We would like to acknowledge Norman Tran for thoughtful discussions. We would also like to acknowledge CHESS staff, specifically Irina Kriksunov, Bill Miller, and David Schuller.

## Author contributions

Conceptualization: MJM, TH, RET

Methodology: MJM, SB, RET

Investigation: MJM, SB, RET

Expression/Purification: MJM, SB

Kinetics: MJM

Crystallography: MJM

Analysis: MJM, RET

Visualization: MJM

Funding acquisition: TH, RET

Project administration: TH, RET

Supervision: TH, RET

Writing – original draft: MJM, RET

Writing – review & editing: MJM, RET, TH

## Data and materials availability

Structural datasets are deposited in the PDB (accession numbers found in **Table S3-5**) and all other data can be made available.

