## Supplemental information for "A structural perspective on the temperature-dependent activity of enzymes"

**This PDF file includes:**

Materials and Methods S1.1-5
Figs. S1-7
Tables S1-6
Data S1-4
References

### S.1 Materials and Methods

#### S.1.1 Materials

Nickel-NTA was purchased from Qiagen, and P6-DG desalting resin was purchased from BioRad. HEPES, TCEP and DTT were purchased from Goldbio. GTP and PEP were purchased from ChemImpex. GDP was purchased from Combi-blocks. MDH was purchased from Calzyme. β-sulfopyruvate, oxalate, and phosphoglycolic acid were purchased from Sigma-Aldrich. All other materials were purchased from the highest grade available.

#### S.1.2 Protein Expression/Purification

Rat cytosolic PEPCK was expressed and purified as previously described.[1] Briefly, $HIS_6$-rcPEPCK-SUMO fusion protein was expressed in BL21(DE3) cells for 16-24 hours in auto-induction media.[2] Cells were harvested in 25 mM HEPES-NaOH pH 7.5 + 300 mM NaCl + 10% (w/v) glycerol + 2 mM TCEP, lysed via French press and kept at 4°C throughout the purification. Lysate was clarified by centrifugation and the supernatant was applied to nickel-NTA resin for 1 hour. Following washing of the resin, rcPEPCK was eluted with 25 mM HEPES-NaOH pH 7.5 + 300 mM imidazole + 2 mM TCEP, buffer exchanged by passage through P6-DG column into 25 mM HEPES-NaOH + 2 mM TCEP and left with SUMO protease overnight. Digested samples were applied to a clean nickel-NTA resin for 1 hour, and flow-through containing rcPEPCK alone was collected. After a final buffer exchange into 25 mM HEPES-NaOH pH 7.5 + 10 mM DTT, samples were concentrated to 10 mg/mL and flash-frozen in 30 µL drops by immersion in liquid nitrogen and stored at -80°C.

##### S.1.3 Kinetic measurements

All measurements were performed using a CaryUV100 spectrophotometer with a temperature controller and were completed in duplicate. Samples were equilibrated at each temperature for 5 minutes and temperature stability was evaluated using a thermocouple. The oxidation of NADH to $NAD^+$ was measured at 340 nm via a coupled assay by using malate dehydrogenase.

The assay mix for rcPEPCK was composed of 0.1 M HEPES-NaOH pH 7.5, 10 mM DTT, 300 µM NADH, 1 mM GDP, 4 mM $MgCl_2$, 100 µM $MnCl_2$, 10 mM PEP, and 50 mM $KHCO_3$ (bubbled with dry ice), 10 U of MDH and 2.5 µg of PEPCK. Reactions were initiated by the addition of rcPEPCK and MDH.

Michaelis-Menten data was analyzed using Enzyme Kinetics package from SigmaPlot 11.0. Linear plotting (Eyring/Arrhenius plots) was completed in R or excel.

##### S.1.4 Multi-temperature crystallography

rcPEPCK crystals were grown using the hanging-drop vapor diffusion method. Protein at 10 mg/mL was mixed with mother liquor in 2:4, 3:3, 4:2 µL (protein:mother liquor) ratios. Mother liquor consisted of 0.1 M HEPES-NaOH pH 7.5, 18-28% PEG 3350. Drops were supplemented with 0.5 µL of 100 mM $MnCl_2$ and 10 mM GTP (or 100 mM GDP). Large rod-shaped crystals were soaked in cryoprotectant solution (for consistency with other previously determined structures) consisting of 0.1 M HEPES-NaOH pH 7.5, 30% PEG 3350, 10% PEG 400, 10 mM $MnCl_2$, and 10 mM of both the nucleotide and inhibitor for at least 15 minutes. Crystals were then harvested, moved through NVH oil to remove surface solution and protect the crystal against dehydration, mounted onto a goniometer at the X-ray source, and cooled or warmed to the data collection temperature using a nitrogen gas stream.

X-ray data collection was performed on CHESS beamline ID7B2, using a beam size of roughly 10 μm × 10 μm. Vector scanning was used to collect diffraction images along the long axis of each crystal, limiting the dose delivered to each region. Each crystal was exposed once, at one temperature, to collect a full dataset. Diffraction frames were indexed, integrated, and scaled using DIALS.[3] A $CC_{1/2}$ value of 0.6 was used as a resolution cut-off for consistency, unless there were obvious issues with collection statistics. Data were merged using AIMLESS[4] and molecular replacement was performed using Molrep in CCP4 using PDB ID# 3DT2.[5–7] Model building was completed using real-space refinement in COOT.[8] Automatic refinement was completed using Phenix.[9] Structures were validated using MolProbity servers (collection and model statistics can be found in **Tables S3-5**).[10]

In previous cryocrystallographic rcPEPCK structures, the Ω-loop can exist in two different states. The loop can be well modeled into the observed density in the closed, ordered state as represented by the rcPEPCK-oxalate-GTP complex (PDB 3DT2) or is not modeled largely when in the open, disordered state such as in the holo-form of the enzyme (PDB 2QEW) (**Fig. 1**). Here, the Ω-loop was modelled with 100% occupancy in the closed state regardless of electron density evidence. Occupancy refinements for GTP in the βSP-GTP complex were performed by fixing the B-factor to the average of the protein and using automatic occupancy refinement. In contrast, phosphoglycolic acid (PGA) occupancy refinements were performed by fixing the B-factor of the ligand to the average B-factor of the protein and refining the occupancy in 5% increments until the $F_o$-$F_c$ peaks corresponding to the $2F_o$-$F_c$ density surrounding and within the ligands of interest were minimized (ie. flattest $F_o$-Fc map in region of interest).[11,12] Alternative refinement protocols were performed for both the Ω-loop and PGA (refine both B-factors and occupancy, refine B-factors with fixed occupancy, fixed B-factors and refined occupancy) and yielded consistent electron density map evidence (see **Table S6** for occupancy/B-factors). For PGA occupancy refinements, automated occupancy refinements in phenix.refine were performed but led to more $F_0$-$F_c$ peaks compared to manual refinements.

##### S.1.5 Interpreting changes in Ω-loop electron density

**Fig. 4** shows how the electron density of the Ω-loop evolves with temperature for each of the three inhibitor complexes. For the βSP-GDP complex, there is a loss of density in the ordered, closed-loop conformation at 40°C relative to that at lower temperatures, but the data set resolution at 40°C is significantly worse. The two possible interpretations of this result are that either the apparent increase in loop disorder is a real phenomenon or it is an artifact of the observed decreased resolution of the data.

**Table S2 and Data S3** show the average B-factor for residues 464-474 (Ω-loop) and for the whole protein for the three ligand-bound complexes at each temperature. In real space, the B-factor is related to the mean squared width of the electron density distribution around an atom; it describes the broadening of that electron density relative to that of the (T=0 K, no vibration) atomic electron density given by the atomic form factor. If we make the simplifying assumptions that (1) at our modest data set resolutions, the observed electron density width averaged over the whole protein and for the loop are both large compared with the intrinsic atomic width, (2) the average B-factor for the protein $B_{protein}$ represents a "background" blurring of the electron density (e.g., largely arising from disorder in packing the complexes within the crystal), while $B_{loop}$ is due to contributions from background blurring ($B_{protein}$) and from local disorder of the loop ($B_{loop, disorder}$), and (3) the real space electron density variances ($\propto$B) add, so $B_{loop} = B_{protein} + B_{loop,disorder}$. The contribution to the real space broadening of the electron density in the loop region from loop disorder alone is then given by $(B_{loop} – B_{protein})^{1/2}$. The difference $B_{loop} – B_{protein}$ should thus be a better proxy for loop disorder than $B_{loop}$ or the normalized $B_{loop}/ B_{protein}$. This difference is also given in **Table S2** and represented in **Data S3**.

For the βSP-GTP complex, both the maps in **Fig. 4 and Fig. S5** and the B factor difference (**Table S2**) indicate a small increase in closed conformation loop density / loop order from -20 C to 20 °C and then a loss of density /order at 40 °C.

For the OX-GTP complex, both the maps in **Fig. 4** and the B factor difference (**Table S2**) indicate a small increase in closed conformation loop density / loop order from -20 C to 20 °C and then a decrease of density/order at 40 °C.

For the PGA-GDP complex, both the maps in **Figs. 4** and the B factor difference (**Table S2**) indicate a gradual increase in closed conformation loop density /order with increasing temperature over the whole temperature range (even though resolution decreases continuously.)

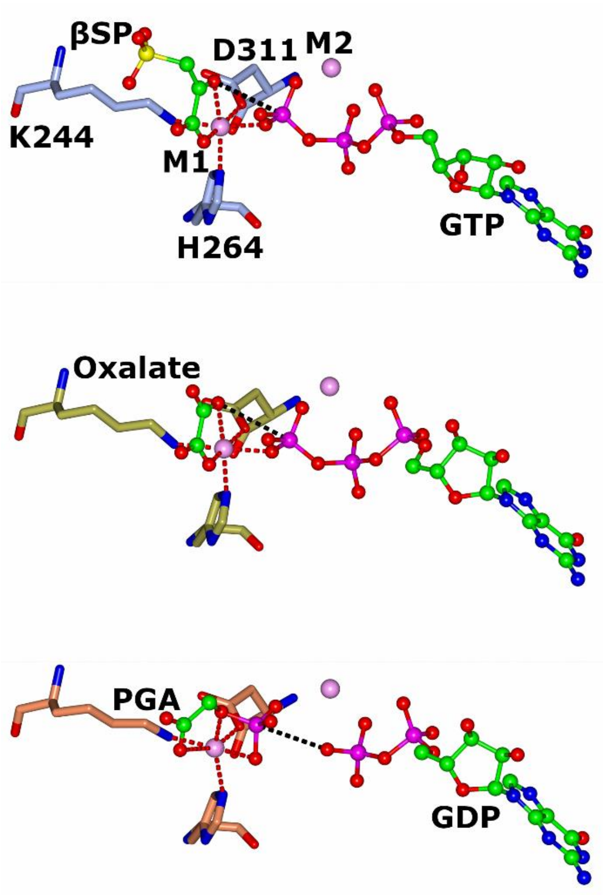


| Complex | Interaction | d (Å) |
|---|---|---|
| **βSP-GTP** | M1 - D311 OD1 | 2.1 |
| | M1 - H264 NE2 | 2.3 |
| | M1 - K244 Nz | 2.3 |
| | M1 - βSP O1 | 2.1 |
| | M1 - βSP O2 | 2.4 |
| | M1 - GTP O2G | 2.2 |
| | βSP O2 - GTP PG | 3.4 |
| **OX-GTP** | M1 - D311 OD1 | 2 |
| | M1 - H264 NE2 | 2.3 |
| | M1 - K244 Nz | 2.3 |
| | M1 - OXL O1 | 2.4 |
| | M1 - OXL O2 | 2.3 |
| | M1 - GTP O2G | 2.3 |
| | βSP O2 - GTP PG | 3.5 |
| **PGA-GDP** | M1 - D311 OD1 | 2.1 |
| | M1 - H264 NE2 | 2.4 |
| | M1 - K244 Nz | 2.2 |
| | M1 - PGA O1 | 3.2 |
| | M1 - PGA O1P | 2.4 |
| | M1 - PGA O2P | 2.2 |
| | PGA P - GTP O2B | 3.4 |

**Fig. S1. Interaction distances of M1 metal and phosphoryl transfer distance for each complex.** Each of the three complexes A) βSP-GTP (ice blue), B) oxalate-GTP (gold), C) PGA-PGA (coral) have mostly conserved interaction distances between active site and ligands with the M1 metal (red dotted line) and distances for phosphoryl acceptor-donor paired (black dotted line)**.** All other atoms are colored by type.

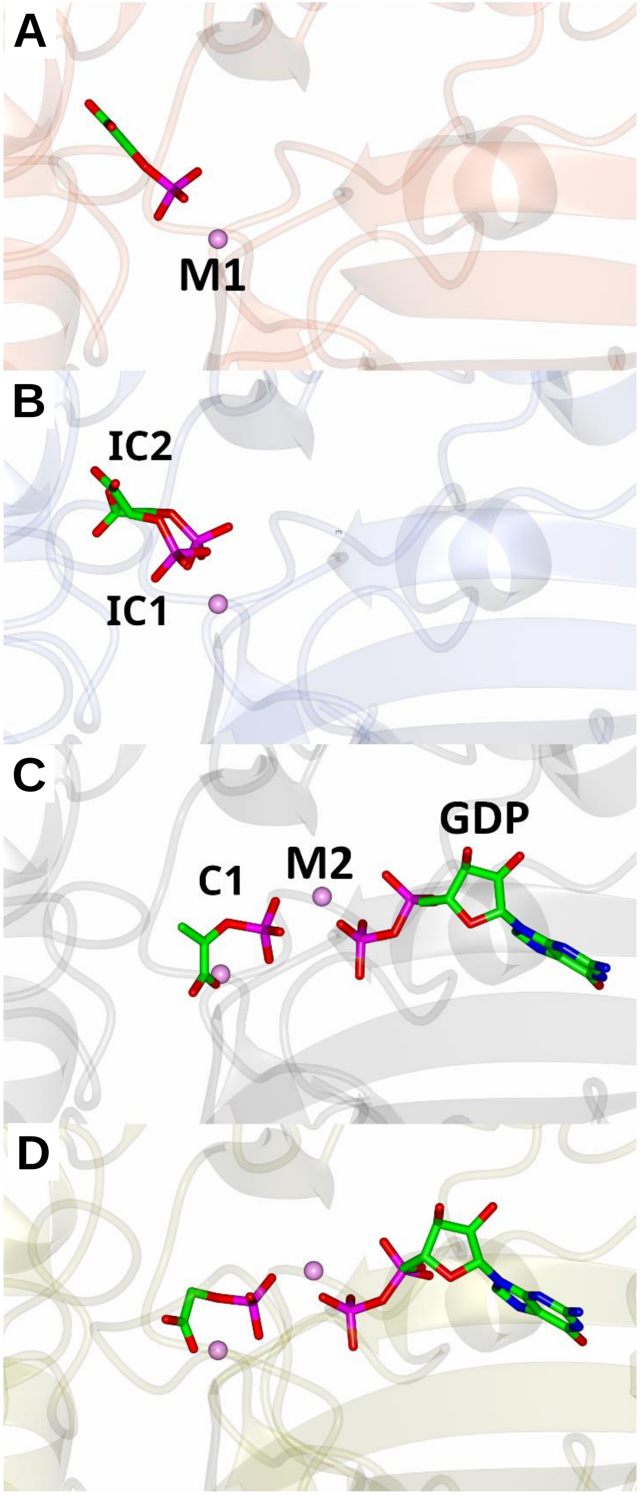


**Fig. S2. Cryogenic structures illustrating variable binding modes of PGA and PEP.** A) PDB 2QZY – PEP bound in outer, incompetent conformation. B) PDB 2RKA – PGA bound in two incompetent conformations. C) PDB 7L3M – PEP bound with GDP in the competent conformation. D) PDB 3DTB – PGA with GDP bound in the competent formation.

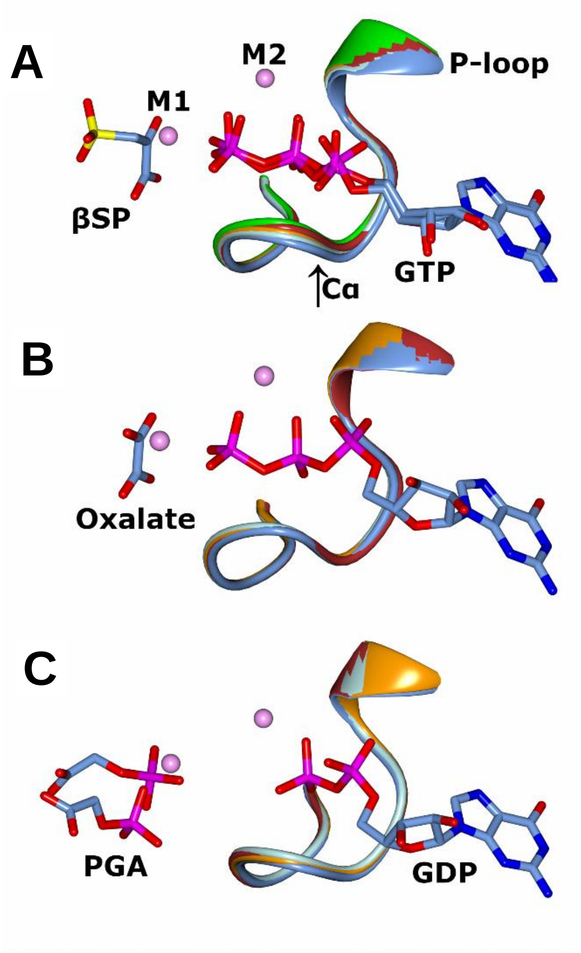


**Fig. S3. P-loop transition between -20°C and 40°C for each complex. A)** Superposition of βSP-GTP complexes showing a shift in the P-loop location and nucleotide positioning toward the metal centers with increasing temperature (-20°C **-** ice blue, 0°C – pale blue, 20°C – orange, 40°C – firebrick red, oxalate-GTP at 20°C – green). When comparing the positions of A287 Cα, it increasingly changes position relative to its position at -20°C (-20°C vs. 0°C – 0.38 Å; -20°C vs. 20°C – 0.63 Å, -20°C vs. 40°C - 0.83 Å). **B)** Superimposition of oxalate-GTP complexes and **C)** superimposition of PGA-GDP complexes showing relatively little change in the P-loop orientation with temperature. Structures aligned based on the whole protein chain."

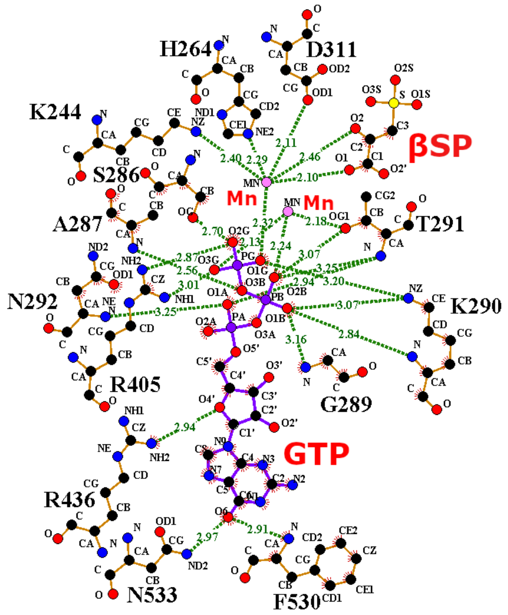


| Interaction | $d_{253}$ (Å) | $d_{313}$ (Å) | Δd |
|---|---|---|---|
| S286 OG - GTP O2G | 3.66 | *2.70 | 0.96 |
| K290 NZ - GTP O1G | 2.75 | 3.20 | -0.45 |
| K290 NZ - GTP O1B | 2.65 | 3.07 | -0.42 |
| K290 N - GTP O1B | 2.98 | 2.84 | 0.14 |
| R405 NH1 - GTP O3G | 3.16 | 3.01 | 0.15 |
| R405 NH2 - GTP O2G | 2.84 | 2.87 | -0.03 |
| A287 N - GTP O3B | 3.27 | 2.56 | 0.71 |
| T291 OG1 - GTP O2B | 3.09 | 3.07 | 0.02 |
| T291 N - GTP O2B | 2.96 | 3.25 | -0.29 |
| T291 N - GTP O1A | 2.96 | 2.94 | 0.02 |
| G289 N - GTP O1B | 3.01 | 3.16 | -0.15 |
| G289 N - GTP O5' | 3.25 | 3.87 | -0.62 |
| R436 NH2 - GTP O4' | 3.49 | 2.94 | 0.55 |
| N292 N- GTP O1A | 5.24 | 3.25 | 1.99 |
| F530 N - GTP O6 | 2.83 | 2.91 | -0.08 |
| N533 ND2 - GTP O6 | 2.94 | 2.97 | -0.03 |

***Uncertainty in position**

**Fig. S4. LigPlot+ PEPCK-GTP interactions at 313 K and changes upon warming.** Ligand interaction map indicating some of the H-bonds of the βSP-GTP complex at 313 K. Atoms are coloured by type: nitrogen – blue, oxygen – red, carbon – black, phosphorous – purple, sulfur – yellow, manganese – pink. Table indicates the changes in interactions upon warming from 253 – 313 K. Heatmap is colored from -1.99 Å (blue) to 0 (white) +1.99 Å **(**red**).**

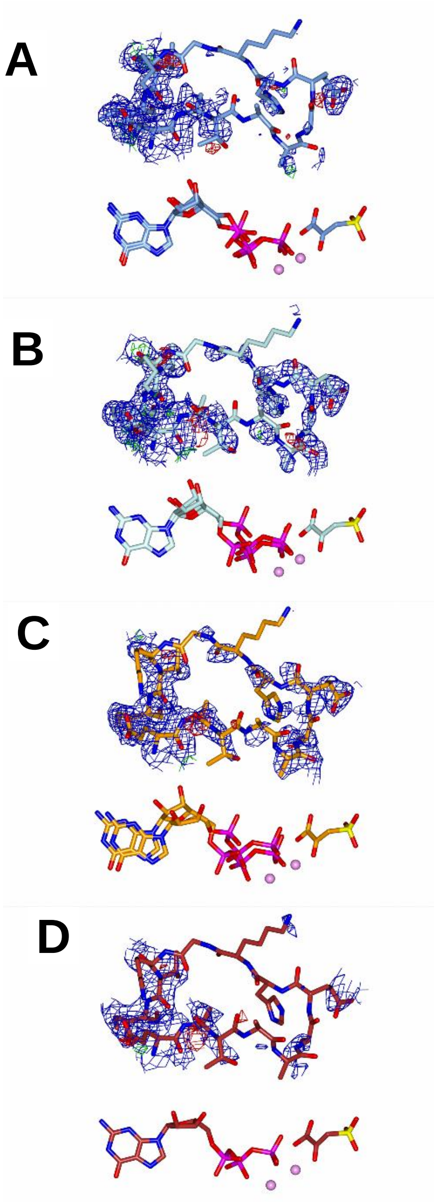


**Fig. S5. Lower contoured electron density of Ω-loop in βSP-GTP complex.** A) -20°C, B) 0°C, C) 20°C, D) 40°C. $2F_o$-$F_c$ electron density map is contoured to 0.7 σ, $F_o$-$F_c$ maps are contoured to 3 σ. Lower threshold maps more clearly show ordering of the Ω-loop from -20 - 20°C and the onset of disorder at 40°C. There is similar amounts of electron density at 0 and 20°C.

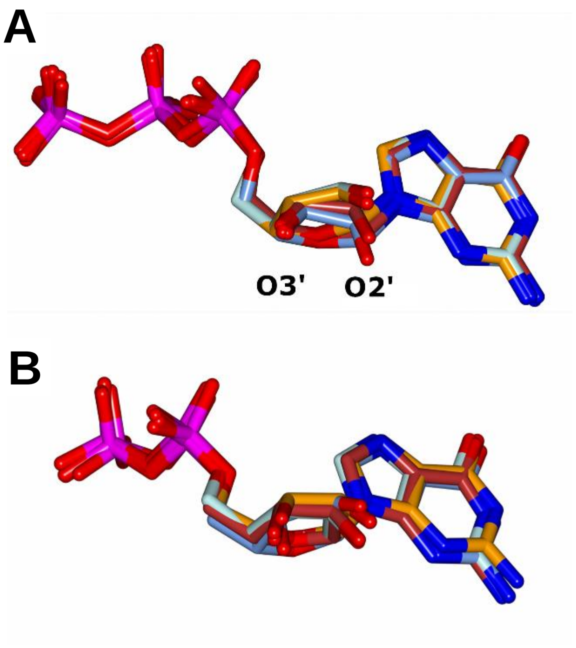


**Fig. S6. Minor conformation changes in nucleotide for intermediate and product complex.** A) Superimposition of structures of GTP in the oxalate-GTP complexes between -20°C and 40°C. B) Superimposition of GDP in the PGA-GDP complex between -20°C and 40°C. These alignments show a small rotational change of the ribose moiety.

**Oxalate-GTP** *In all cases, the B-factor for O2’ is the largest B-factor for any atom in the nucleotide*

| Temperature (°C) | O2’ B-factor ($Å^2$) | GTP average B-factor ($Å^2$) | O2’/GTP |
|---|---|---|---|
| -20 | 63.1 | 48.4 | 1.30 |
| 0 | 49.1 | 39.3 | 1.25 |
| 20 | 47.3 | 35.6 | 1.32 |
| 40 | 65.8 | 52.9 | 1.24 |

**PGA-GDP** *The B-factor for O3’ is the largest B-factor for any atom in the nucleotide for only at -20°C and 20 °C. For 0 °C and 40 °C they are ranked 2$^{nd}$ and 4$^{th}$ respectively but the atoms with larger B-factors are all on the ribose moiety*

| Temperature (°C) | O3’ B-factor ($Å^2$) | GTP average B-factor ($Å^2$) | O3’/GTP |
|---|---|---|---|
| -20 | 55.5 | 45.3 | 1.22 |
| 0 | 49.7 | 39.9 | 1.24 |
| 20 | 72.6 | 58.8 | 1.29 |
| 40 | 77.4 | 67.9 | 1.14 |

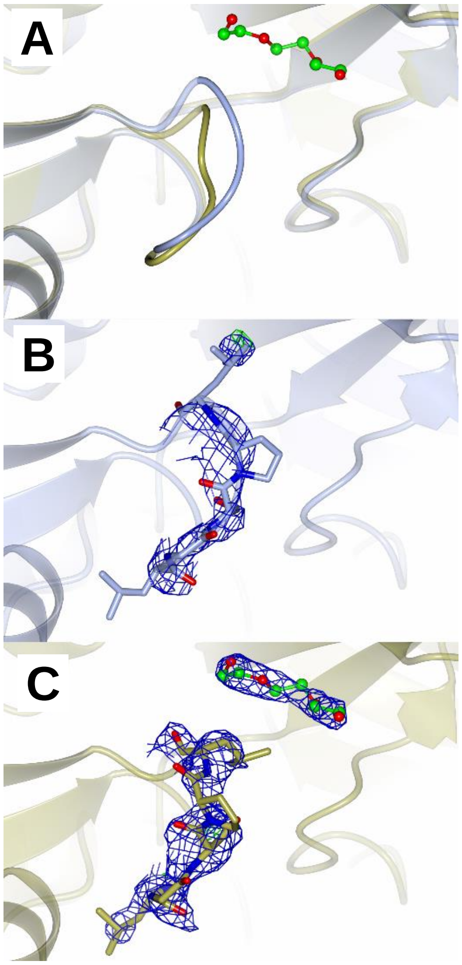


**Fig. S7. PEG binding site.** PEG (green) is shown in its binding site. A) There are two known conformations of loop 149-153 depending on the ligands present at the active site. B) When rcPEPCK is in complex with BSP-GTP or PGA-GDP (shown in ice blue) loop 149-154 adopts a conformation which occludes PEG binding. C) When rcPEPCK is in complex with oxalate-GTP (gold) loop 149-154 adopts an open conformation conducive of PEG binding. All other atoms are colored by type.

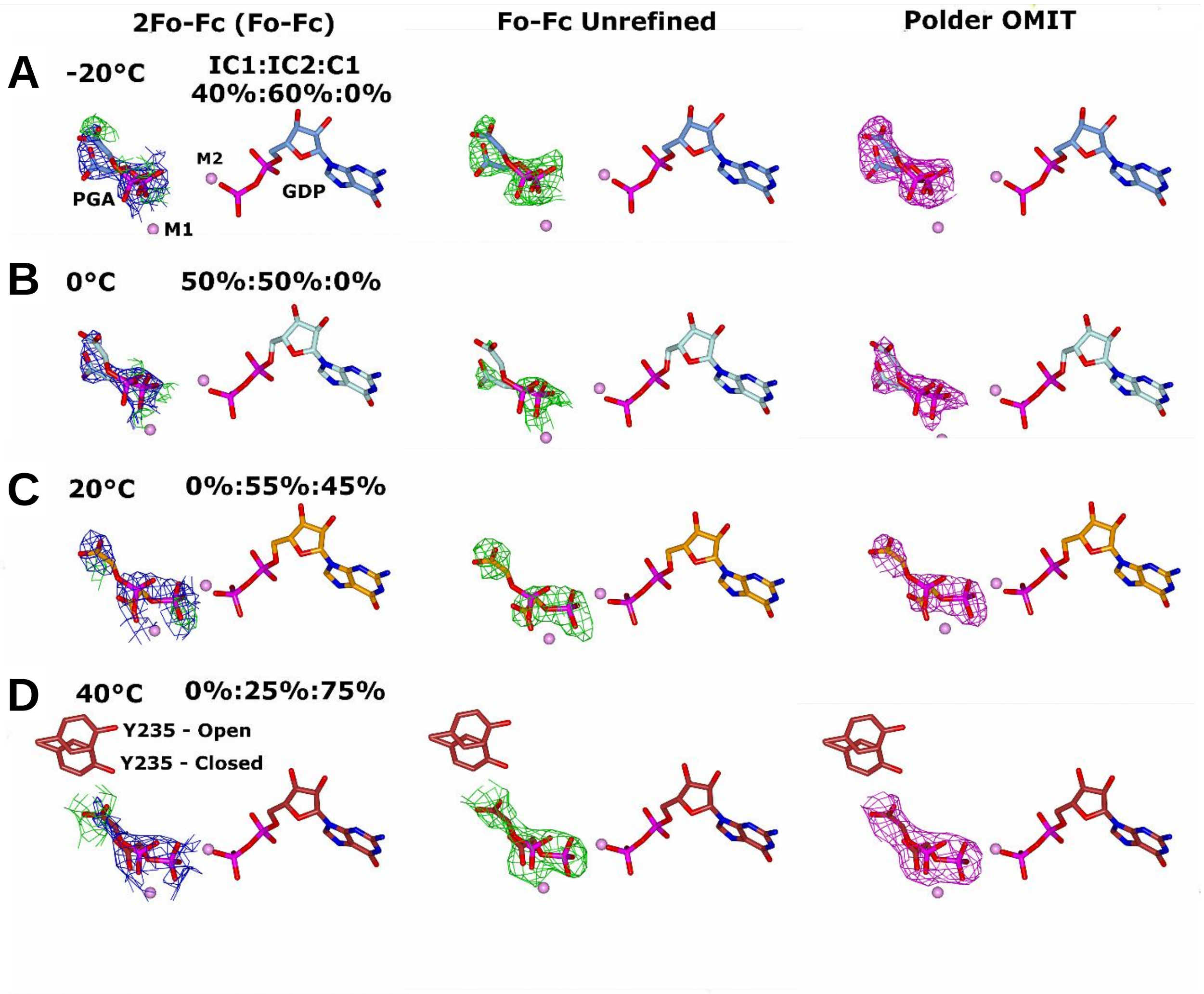


**Fig. S8. Electron density maps illustrating the shift in the conformational position of PGA toward the competent conformation (C1) as with increasing temperature.** Adaptation of Figure 4 showing additional electron density maps. A) -20°C, B) 0°C, C) 20°C, D) 40°C PGA-GDP complex. The left panel shows $2F_o$-$F_c$ (blue - 1σ RMSD) and $F_o$-$F_c$ (green and red - 3σ) after PGA is modelled and refined. Center panel shows $F_o$-$F_c$ (green and red – 3σ) prior to PGA modeling and refinement. The right panel shows the Polder OMIT map (magenta - 5σ) focused on PGA.

**Table S1.**

Temperature dependence of $k_{cat}/K_M$ for rcPEPCK in the PEP + GDP + $CO_2$ → OAA + GTP catalyzed reaction. Errors are not standard error of replicates but rather derived by computed $k_{cat}$ and $K_M$ error as percentage, then propagating these errors.

| **Temperature (°C)** | **$k_{cat}/K_M$ ($M^{-1}s^{-1}$)** |
|---|---|
| 15 | $5.0 \times 10^4 \pm 3.7 \times 10^3$ |
| 25 | $6.5 \times 10^4 \pm 7.7 \times 10^3$ |
| 37 | $2.2 \times 10^5 \pm 2.2 \times 10^4$ |
| 55 | $2.8 \times 10^5 \pm 3.8 \times 10^4$ |

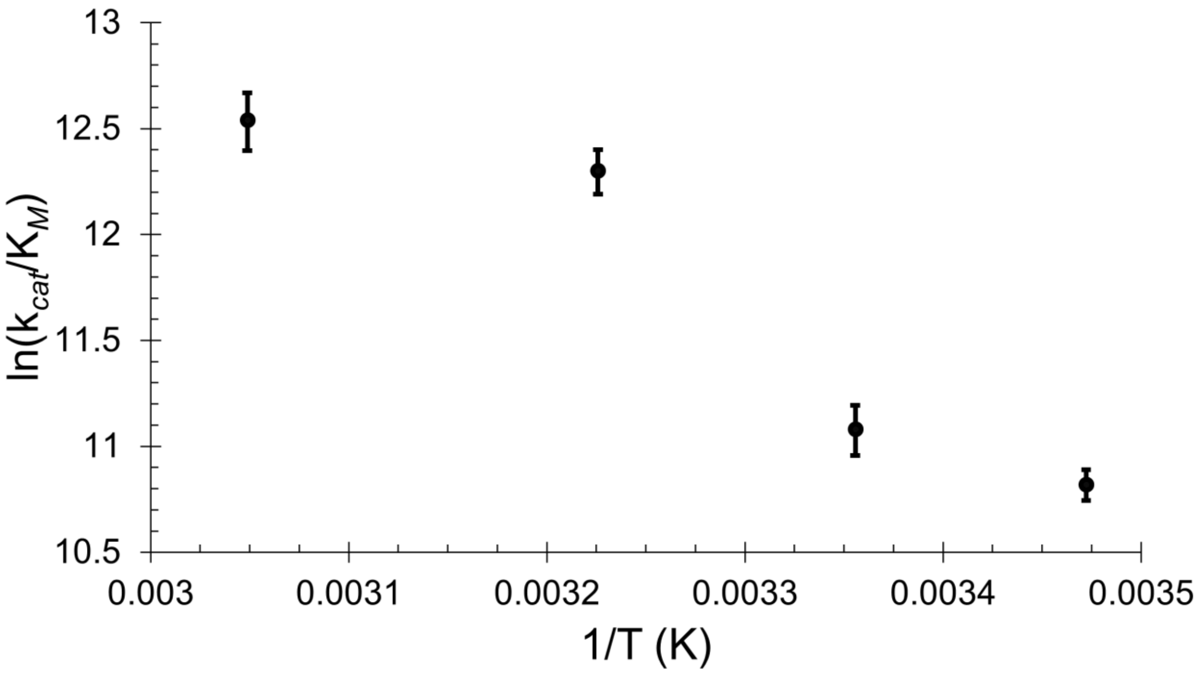

**Table S2.** Average B-factors for the Ω-loop (residues 464-474) and for the entire protein and corresponding data set resolution for each ligand-bound complex at each temperature. The difference $B_{loop} – B_{protein}$ provides a measure of loop disorder that accounts for differences in data set resolution (**SI Sec. S1.5**)

| Temp (°C) | $B_{loop}$ (Å$^2$) | $B_{protein}$ (Å$^2$) | $B_{loop}$ - $B_{protein}$ (Å$^2$) | Resolution (Å) |
|---|---|---|---|---|
| **βSP-GTP** | | | | |
| -20 | 102 | 33.6 | 68.4 | 1.86 |
| 0 | 90.3 | 33.9 | 56.4 | 1.74 |
| 20 | 83.4 | 34.9 | 48.5 | 1.94 |
| 40 | 125 | 55.5 | 69.5 | 2.41 |
| **OX-GTP** | | | | |
| -20 | 84.6 | 55.5 | 29.1 | 2.04 |
| 0 | 72.7 | 49 | 23.7 | 1.95 |
| 20 | 72.8 | 46.4 | 26.4 | 2 |
| 40 | 106 | 64.1 | 41.9 | 2.12 |
| **PGA-GDP** | | | | |
| -20 | 121 | 53.7 | 67.3 | 1.91 |
| 0 | 131 | 65.6 | 65.4 | 2.2 |
| 20 | 135 | 80.7 | 54.3 | 2.42 |
| 40 | 129 | 76.5 | 52.5 | 2.53 |

**Table S3.**

Crystallographic data for the rcPEPCK-βSP-GTP complex.

| | βSP-GTP complex at -20°C | βSP-GTP complex at 0°C | βSP-GTP complex at 20°C | βSP-GTP complex at 40°C |
|---|---|---|---|---|
| **PDB ID** | 8SYQ | 8SYR | 8SYS | 8SYT |
| **Wavelength (Å)** | 1.1271 Å | 1.1271 Å | 1.1271 Å | 1.1271 Å |
| **Resolution range (Å)** | 41.8 - 1.86 (1.93 - 1.86) | 41.8 - 1.74 (1.81 - 1.74) | 51.0 - 1.94 (2.01 - 1.94) | 56.7 - 2.41 (2.49 - 2.41) |
| **Space group** | P 1 21 1 | P 1 21 1 | P 1 21 1 | P 1 21 1 |
| **Unit cell (Å, °)** | 44.85 119.77° 60.55 90° 111.12 90° | 44.85 119.83° 60.57 90° 111.21 90° | 44.82 119.74° 60.55 90° 111.39 90° | 44.90 120.20° 60.98 90° 111.49 90° |
| **Total reflections** | 331627 (32181) | 385769 (25594) | 297884 (29952) | 78319 (6977) |
| **Unique reflections** | 49155 (4808) | 58489 (4850) | 43110 (4235) | 23367 (2303) |
| **Multiplicity** | 6.7 (6.7) | 6.6 (5.3) | 6.9 (7.1) | 3.4 (3.0) |
| **Completeness (%)** | 98.4 (97.3) | 96.4 (79.5) | 98.1 (96.7) | 99.5 (96.8) |
| **Mean I/sigma(I)** | 7.05 (0.90) | 9.79 (0.97) | 3.39 (0.48) | 3.94 (0.83) |
| **Wilson B-factor (Å$^2$)** | 24.2 | 25.1 | 24.8 | 37.9 |
| **R-merge** | 0.171 (1.452) | 0.111 (0.923) | 0.15 (1.379) | 0.136 (0.833) |
| **R-meas** | 0.186 (1.575) | 0.120 (1.024) | 0.162 (1.487) | 0.162 (1.012) |
| **R-pim** | 0.071 (0.605) | 0.045 (0.430) | 0.061 (0.554) | 0.086 (0.568) |
| **CC1/2** | 0.991 (0.648) | 0.992 (0.694) | 0.995 (0.657) | 0.982 (0.643) |
| **Reflections used in refinement** | 49131 (4803) | 58323 (4775) | 43076 (4231) | 23317 (2271) |
| **Reflections used for R-free** | 2432 (241) | 2934 (225) | 2190 (226) | 1082 (94) |
| **R-work** | 0.172 (0.291) | 0.163 (0.262) | 0.167 (0.269) | 0.196 (0.267) |
| **R-free** | 0.201 (0.322) | 0.190 (0.268) | 0.198 (0.293) | 0.224 (0.279) |
| **Number of non-hydrogen atoms** | 5316 | 5259 | 5198 | 5032 |
| **macromolecules** | 4895 | 4901 | 4901 | 4903 |
| **ligands** | 77 | 77 | 77 | 45 |
| **solvent** | 344 | 281 | 220 | 84 |
| **Protein residues** | 620 | 620 | 620 | 620 |
| **RMS(bonds) (Å)** | 0.006 | 0.006 | 0.008 | 0.002 |
| **RMS(angles) (°)** | 0.89 | 0.93 | 0.98 | 0.51 |
| **Ramachandran favored (%)** | 96.9 | 97.1 | 96.93 | 95.3 |
| **Ramachandran allowed (%)** | 3.07 | 2.91 | 2.75 | 4.37 |
| **Ramachandran outliers (%)** | 0 | 0 | 0.32 | 0.32 |
| **Rotamer outliers (%)** | 1.15 | 0.58 | 0.96 | 0.38 |
| **Clashscore** | 2.13 | 1.83 | 3.96 | 2.55 |
| **Average B-factor (Å$^2$)** | 33.7 | 33.6 | 35.2 | 55.5 |
| **macromolecules** | 33.6 | 33.52 | 35.2 | 55.8 |
| **ligands** | 31.8 | 27.4 | 32.5 | 38.7 |
| **solvent** | 35.3 | 37.3 | 35.1 | 42.8 |
| **Number of TLS groups** | 5 | 4 | 4 | 6 |

**Table S4.**

Crystallographic data table for the rcPEPCK-oxalate-GTP complex.

| | **Oxalate-GTP complex at -20°C** | **Oxalate-GTP complex at 0°C** | **Oxalate-GTP complex at 20°C** | **Oxalate-GTP complex at 40°C** |
|---|---|---|---|---|
| **PDB ID** | 8SYU | 8SYV | 8SYW | 8SYX |
| **Wavelength (Å)** | 1.1271 Å | 1.1271 Å | 1.1271 Å | 1.1271 Å |
| **Resolution range (Å)** | 56.5 - 2.04<br>(2.11 - 2.04) | 42.0 - 1.95<br>(2.02 - 1.95) | 56.5 - 2.00<br>(2.07 - 2.00) | 56.9 -2.12<br>(2.20 -2.12) |
| **Space group** | P 1 21 1 | P 1 21 1 | P 1 21 1 | P 1 21 1 |
| **Unit cell (Å,°)** | 45.22 120.16°<br>60.64 90° 111.37 90° | 45.28 120.17°<br>60.82 90° 111.84 90° | 45.32 119.83°<br>60.83 90° 111.66 90° | 45.30 120.16°<br>61.12 90° 111.47 90° |
| **Total reflections** | 195478 (10105) | 294402 (29278) | 273012 (26988) | 209071 (9714) |
| **Unique reflections** | 37226 (3435) | 43860 (4330) | 40575 (4003) | 33964 (3333) |
| **Multiplicity** | 6.9 ( 7.1) | 6.7 (6.8) | 6.7 (6.7) | 6.9 (6.5) |
| **Completeness (%)** | 93.7 (65.3) | 99.8 (99.8) | 99.8 (98.4) | 88.2 (21.8) |
| **Mean I/sigma(I)** | 1.9 (0.3) | 6.47 (0.65) | 2.87 (0.44) | 4.9 (0.7) |
| **Wilson B-factor ($Å^2$)** | 41.0 | 35.4 | 36.5 | 49.0 |
| **R-merge** | 0.412 (2.101) | 0.104 (1.339) | 0.244 (1.581) | 0.297 (1.481) |
| **R-meas** | 0.448 (2.270) | 0.113 (1.451) | 0.265 (1.713) | 0.321 (1.611) |
| **R-pim** | 0.174 (0.852) | 0.043 (0.555) | 0.101 (0.655) | 0.122 (0.628) |
| **CC1/2** | 0.892 (0.649) | 0.997 (0.676) | 0.976 (0.663) | 0.968 (0.607) |
| **Reflections used in refinement** | 35852 (2478) | 43831 (4327) | 40573 (4004) | 30375 (747) |
| **Reflections used for R-free** | 1667 (134) | 2244 (219) | 2032 (227) | 1412 (41) |
| **R-work** | 0.207 (0.386) | 0.172 (0.295) | 0.179 (0.286) | 0.202 (0.393) |
| **R-free** | 0.264 (0.417) | 0.205 (0.339) | 0.213 (0.330) | 0.227 (0.380) |
| **Number of non-hydrogen atoms** | 5016 | 5098 | 5129 | 5155 |
| **macromolecules** | 4878 | 4892 | 4908 | 4902 |
| **ligands** | 51 | 51 | 51 | 51 |
| **solvent** | 87 | 155 | 170 | 202 |
| **Protein residues** | 620 | 620 | 620 | 620 |
| **RMS(bonds) (Å)** | 0.002 | 0.007 | 0.002 | 0.002 |
| **RMS(angles) (°)** | 0.54 | 0.9 | 0.57 | 0.44 |
| **Ramachandran favored (%)** | 96.4 | 96.4 | 96.8 | 95.8 |
| **Ramachandran allowed (%)** | 3.40 | 3.56 | 3.24 | 4.05 |
| **Ramachandran outliers (%)** | 0.16 | 0 | 0 | 0.16 |
| **Rotamer outliers (%)** | 0.39 | 0.96 | 0.38 | 0 |
| **Clashscore** | 2.76 | 2.86 | 3.15 | 3.05 |
| **Average B-factor ($Å^2$)** | 55.3 | 48.8 | 46.2 | 64.0 |
| **macromolecules** | 55.5 | 49.0 | 46.4 | 64.1 |
| **ligands** | 51.8 | 42.9 | 39.3 | 57.0 |
| **solvent** | 48.0 | 44.2 | 42.3 | 61.2 |
| **Number of TLS groups** | 3 | 3 | 3 | 3 |

**Table S5.**

Crystallographic data table for the rcPEPCK-PGA-GDP complex.

| | **PGA-GDP complex at -20°C** | **PGA-GDP complex at 0°C** | **PGA-GDP complex at 20°C** | **PGA-GDP complex at 40°C** |
|---|---|---|---|---|
| **PDB ID** | 8SYY | 8SYZ | 8SZ0 | 8SZ1 |
| **Wavelength (Å)** | 1.1271 Å | 1.1271 Å | 1.1271 Å | 1.1271 Å |
| **Resolution range (Å)** | 55.9 - 1.91 (1.98 - 1.91) | 51.0 - 2.20 (2.28 -2.20) | 56.6 - 2.42 (2.51 - 2.42) | 57.6 - 2.53 (2.62 - 2.53) |
| **Space group** | P 1 21 1 | P 1 21 1 | P 1 21 1 | P 1 21 1 |
| **Unit cell (Å °)** | 44.71 119.79° 60.07 90° 111.48 90° | 45.13 120.19° 60.55 90° 111.39 90° | 45.42 120.89° 60.83 90° 111.52 90° | 45.89 120.17° 61.41 90° 110.19 90° |
| **Total reflections** | 286852 (28194) | 198252 (18510) | 145779 (15160) | 138551 (7047) |
| **Unique reflections** | 43809 (4167) | 30359 (3004) | 23168 (2291) | 20927 (1036) |
| **Multiplicity** | 6.5 (6.8) | 6.5 (6.2) | 6.3 (6.6) | 6.6 (6.8) |
| **Completeness (%)** | 94.5 (74.4) | 97.2 (79.5) | 86.1 (17.6) | 100 (100) |
| **Mean I/sigma(I)** | 9.69 (0.30) | 5.68 (0.35) | 4.19 (0.17) | 4.9 (0.3) |
| **Wilson B-factor ($Å^2$)** | 40.7 | 49.3 | 63.8 | 59.2 |
| **R-merge** | 0.074 (1.282) | 0.130 (1.788) | 0.149 (1.77) | 0.213 (0.947) |
| **R-meas** | 0.080 (1.388) | 0.1414 (1.955) | 0.162 (1.922) | 0.213 (0.947) |
| **R-pim** | 0.031 (0.528) | 0.055 (0.780) | 0.064 (0.735) | 0.082 (0.362) |
| **CC1/2** | 0.999 (0.668) | 0.997 (0.657) | 0.996 (0.628) | 0.988 (0.859) |
| **Reflections used in refinement** | 42946 (3370) | 29657 (2427) | 20019 (407) | 20192 (1498) |
| **Reflections used for R-free** | 2176 (172) | 1412 (127) | 961 (21) | 990 (73) |
| **R-work** | 0.191 (0.421) | 0.235 (0.385) | 0.231 (0.395) | 0.214 (0.387) |
| **R-free** | 0.226 (0.421) | 0.273 (0.427) | 0.278 (0.388) | 0.249 (0.370) |
| **Number of non-hydrogen atoms** | 5106 | 5032 | 4948 | 4979 |
| **macromolecules** | 4939 | 4931 | 4915 | 4943 |
| **ligands** | 21 | 21 | 21 | 21 |
| **solvent** | 146 | 80 | 12 | 15 |
| **Protein residues** | 620 | 620 | 620 | 620 |
| **RMS(bonds) (Å)** | 0.004 | 0.002 | 0.002 | 0.002 |
| **RMS(angles) (°)** | 0.73 | 0.47 | 0.44 | 0.49 |
| **Ramachandran favored (%)** | 96.9 | 94.7 | 94.3 | 94.3 |
| **Ramachandran allowed (%)** | 2.91 | 5.34 | 5.50 | 5.50 |
| **Ramachandran outliers (%)** | 0.16 | 0.16 | 0.16 | 0.16 |
| **Rotamer outliers (%)** | 0.38 | 0.19 | 0.19 | 0.38 |
| **Clashscore** | 3.66 | 4.69 | 3.78 | 3.35 |
| **Average B-factor ($Å^2$)** | 53.4 | 64.9 | 80.5 | 76.4 |
| **macromolecules** | 53.7 | 65.6 | 80.7 | 76.5 |
| **ligands** | 54.2 | 50.4 | 79.3 | 73.3 |
| **solvent** | 47.3 | 34.8 | 55.6 | 60.5 |
| **Number of TLS groups** | 4 | 4 | 9 | 4 |

**Table S6.** Occupancies and B-factors for PGA after alternative refinement strategies

| Temp (°C) | IC1:IC2:C1 Occupancy | B-Factor (IC1) | B-Factor (IC2) | B-Factor (C1) | B-Factor (macromolecule) | Protocol |
|---|---|---|---|---|---|---|
| -20 | 55:45:0 | 42.7 | 49.3 | 0 | 53.2 | Refine both |
| | 40:60:0 | 52.3 | 52.3 | 0 | 53.7 | Fix B, refine occu. (man.) |
| | 40:60:0 | 51.7 | 51.7 | 0 | 53.7 | Fix B, refine occu. (auto.) |
| 0 | 50:50:0 | 52.6 | 57.7 | 0 | 63.7 | Refine both |
| | 50:50:0 | 65.3 | 65.3 | 0 | 65.6 | Fix B, refine occu. (man.) |
| | 39:61:0 | 48.1 | 48.1 | 0 | 65.6 | Fix B, refine occu. (auto.) |
| 20 | 0:40:60 | 0 | 74.2 | 70.3 | 61.0 | Refine both |
| | 0:55:45 | 0 | 77.0 | 77.0 | 80.7 | Fix B, refine occu. (man.) |
| | 0:56:44 | 0 | 78.3 | 78.3 | 80.7 | Fix B, refine occu. (auto.) |
| 40 | 0:15:85 | 0 | 66.7 | 69.6 | 69.4 | Refine both |
| | 0:25:75 | 0 | 72.4 | 72.4 | 76.5 | Fix B, refine occu. (man.) |
| | 0:43:57 | 0 | 72.9 | 72.9 | 76.5 | Fix B, refine occu. (auto.) |

**Data S1.**

Raw kinetic data for Eyring plot.

| | No Viscogen | | Viscogen | |
|---|---|---|---|---|
| **Temperature (K)** | **ln($k_{cat}$/T)** | **ln($k_{cat}$/T)** | **ln($k_{cat}$/T)** | **ln($k_{cat}$/T)** |
| 280 | | | -5.09 | -4.84 |
| 281 | -5.04 | -4.98 | | |
| 283 | -4.63 | -4.58 | | |
| 284 | | | -4.34 | -4.36 |
| 288 | -3.83 | -3.82 | | |
| 289 | | | -3.47 | -3.47 |
| 293 | -2.97 | -2.98 | -2.82 | -2.84 |
| 298 | -2.36 | -2.39 | -2.22 | -2.34 |
| 303 | -1.85 | -1.99 | -1.74 | -1.86 |
| 307 | -1.39 | -1.42 | -1.21 | -1.45 |
| 312 | -1.13 | -1.03 | | |
| 313 | | | -0.97 | -0.99 |
| 317 | -0.65 | -0.72 | | |
| 318 | | | -0.65 | -0.73 |
| 322 | -0.34 | -0.41 | | |
| 323 | | | -0.22 | -0.81 |
| 327 | -0.06 | -0.32 | | |
| 328 | | | -0.73 | -0.88 |
| 333 | 0.12 | 0.11 | -0.65 | -0.50 |
| 336 | -0.08 | 0.07 | | |

**Data S2.**

Evidence for linear region of PEP→OAA Eyring plot

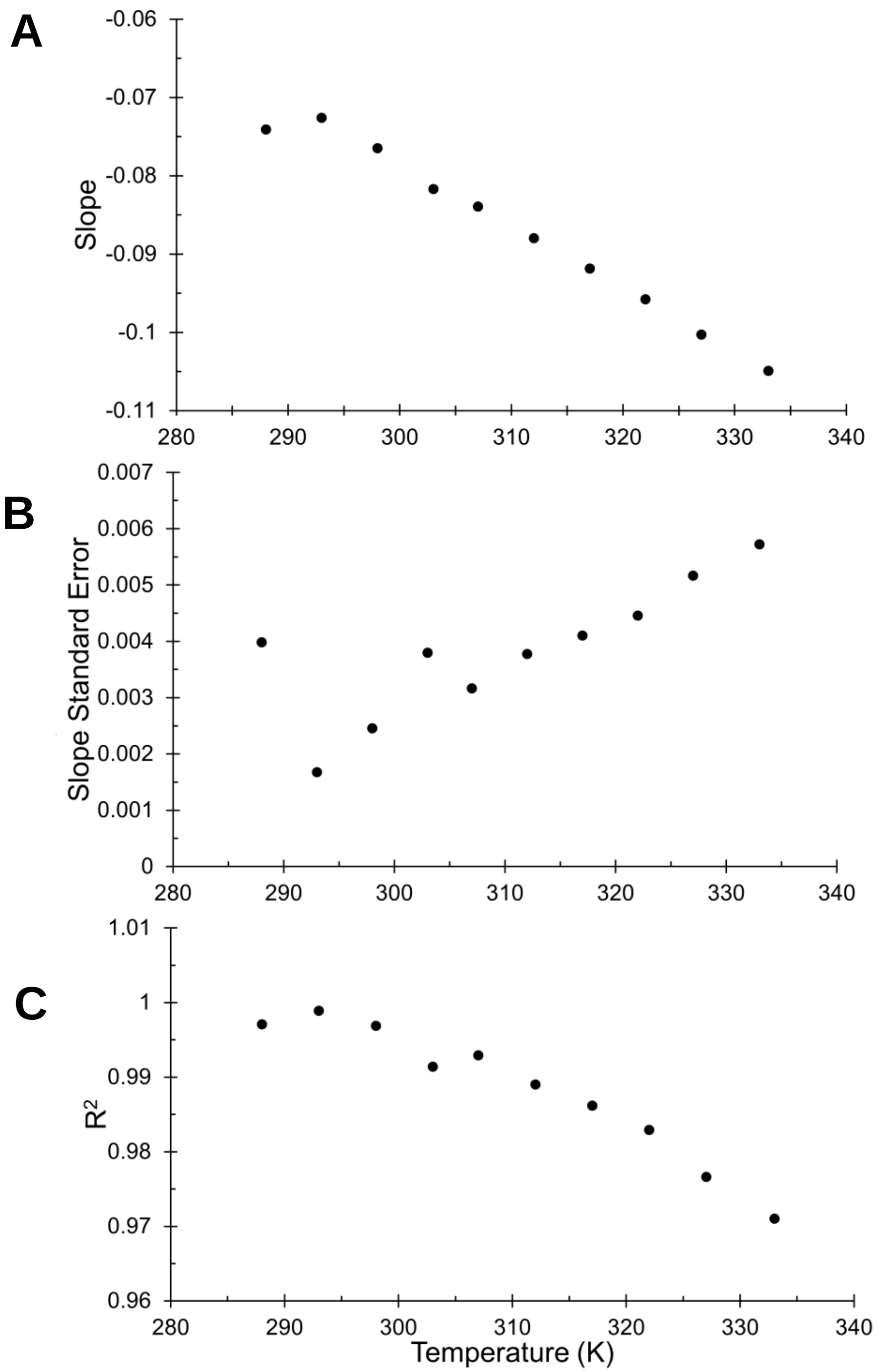


Plots were produced by iteratively calculating A) slope, B) standard error of slope, and C) $R^2$ of data's linear fit after extending the data range. Temperature indicates the upper limit of the temperature range (281 K to displayed temperature).

**Data S3.**

**Normalized B-factor for each complex.**

A) βSP-GTP complex B-factor$_{residue(res)}$ - B-factor$_{macromolecule(MM)}$, B) βSP-GTP complex B-factor$_{res}$/B-factor$_{MM}$, C) OX-GTP complex B-factor$_{res}$ - B-factor$_{MM}$, D) OX-GTP complex B-factor$_{res}$/B-factor$_{MM}$, E) PGA-GDP complex B-factor$_{res}$ - B-factor$_{(MM}$, F) PGA-GDP complex B-factor$_{res}$/B-factor$_{MM.}$ Dotted lines indicate regions that are referenced in main text.

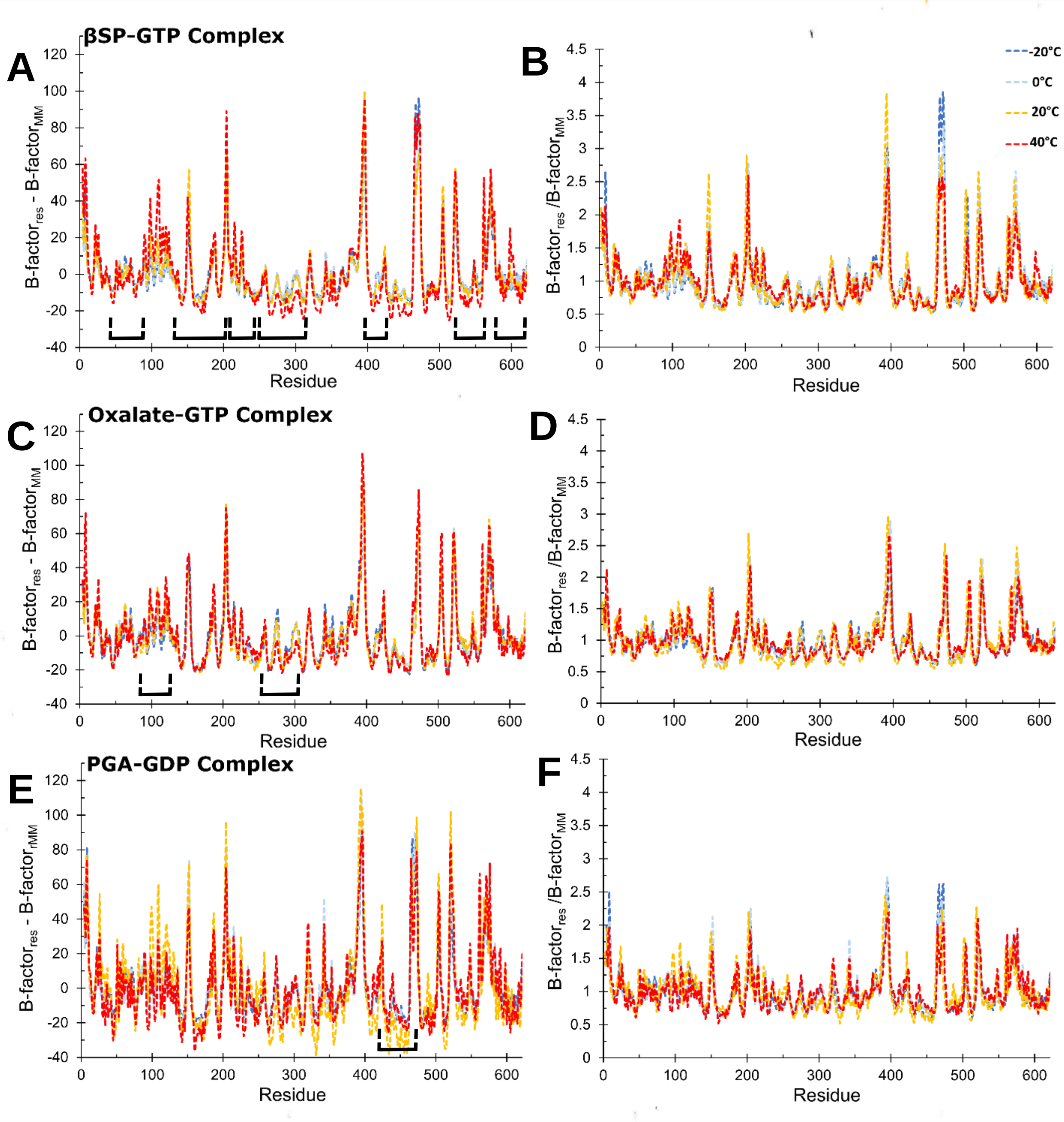

**Data S4.**

Average B-factor (A) and resolution (B) of each complex and temperature.

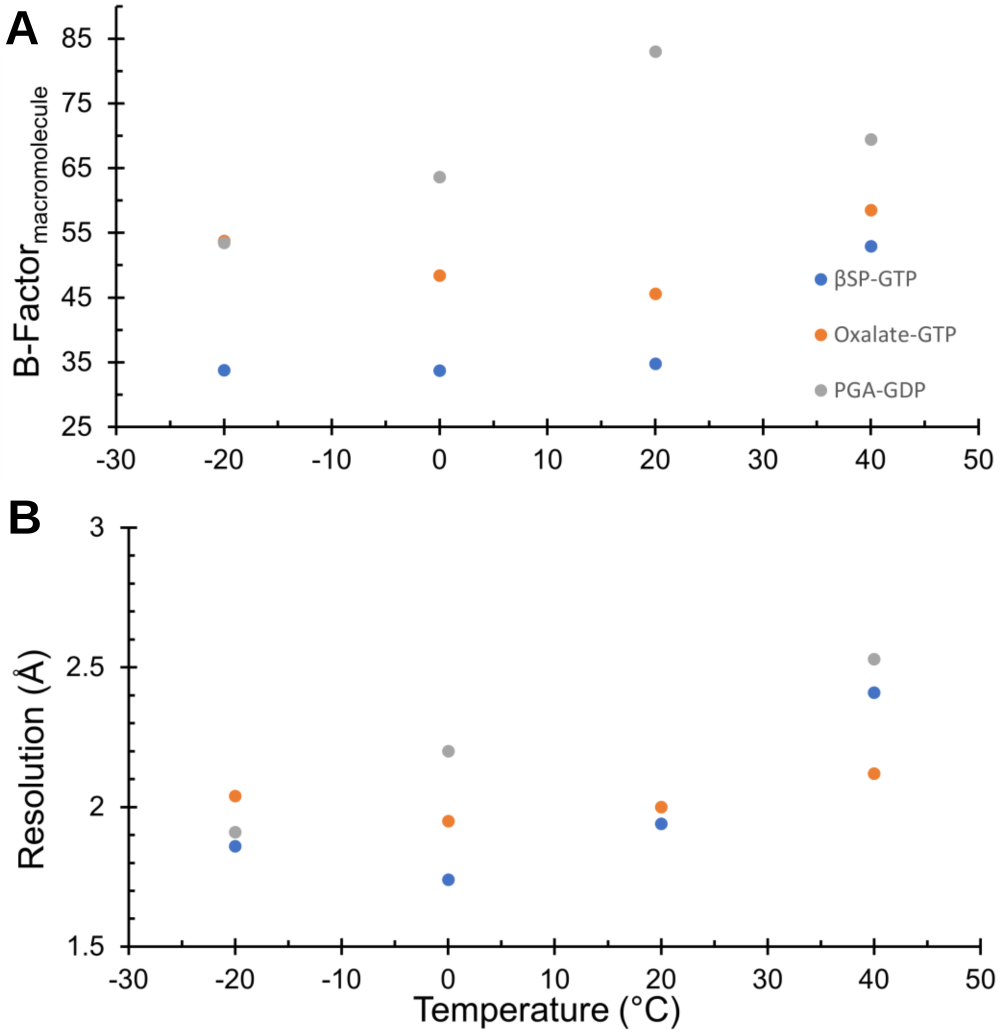
